# Cytoplasmic FKBP7 reflux sustains NFE2L1 levels in an adaptive response to chemotherapy in prostate cancer cells

**DOI:** 10.64898/2026.01.18.700172

**Authors:** Luce Dreno, Farès Ousalem, Anilo Albornoz-Loaiza, Besimé Celik, Carolina Tiraboschi, Giuseppina Claps, Nader Al Nakouzi, Daniel Compagno, Thibault Dayris, Marine Aglave, Yohann Loriot, Nadine Assrir, Ewen Lescop, Anne Chauchereau

## Abstract

Understanding how cancer cells adapt to chemotherapy is essential for overcoming drug resistance. Here, we identify a mechanism of taxane resistance in prostate cancer mediated by FKBP7, a poorly characterized *cis-trans* prolyl isomerase resident in the endoplasmic reticulum (ER). Using cell fractionation, we show that FKBP7 undergoes proteasome-independent retrotranslocation from the ER to the cytosol. This cytosolic pool is increased in docetaxel-resistant cells, which have also developed an enhanced capacity to counteract chemotherapy-induced oxidative stress. In the cytosol, FKBP7 associates with the translation machinery, interacting with multiple regions of the scaffolding protein eIF4G1, including its C-terminal HEAT3 domain, as well as with the 40S ribosomal subunit and polysomes. Polysome profiling and RNA sequencing after FKBP7 silencing revealed that FKBP7 loss leads to translation buffering of a large set of mRNAs, many of which are targets of the transcription factor NFE2L1, a master regulator of the oxidative stress response. Consistently, FKBP7 sustained NFE2L1 protein levels, and both proteins were required for the survival of docetaxel-resistant cells. Pharmacological inhibition of NFE2L1 activation suppressed the growth of docetaxel-resistant cells. We further report the first structural characterization of the FKBP7 catalytic domain by NMR, showing that it is a well-folded protein that binds rapamycin and everolimus, but not FK506. Together, these findings uncover a moonlighting function for an ER-resident chaperone in cytosolic translational control and oxidative stress adaptation and identify the FKBP7-NFE2L1 axis as a potential therapeutic target to overcome chemoresistance.

## INTRODUCTION

In Prostate cancer (PCa), taxane-based chemotherapy has shifted from being solely used as a palliative treatment to being combined with therapies targeting the androgen receptor axis, markedly improving survival *(1, 2)*. However, resistance frequently develops, underscoring the need to understand the mechanisms of taxane resistance and to identify strategies to overcome it. Taxane resistance is multifactorial and multiple pathways have been identified in this process *(2)*. Emerging evidence suggests that resistance also involves adaptive changes in response to stress. This adaptation to microenvironmental stress can manifest as transcriptional reprogramming, as observed during epithelial-mesenchymal transition (EMT) *(2)*, endoplasmic reticulum (ER) stress-induced activation of the unfolded protein response (UPR) *(3)*, and adaptation to reactive oxygen species (ROS)-induced stress *(4)*. Cellular adaptation to ER stress is mediated by the UPR, which aims to restore ER homeostasis, and various cancers exhibit increased ER stress in response to chemotherapeutic drugs *(5)*. In addition, it has been shown that ER stress can also trigger the reflux of some correctly folded proteins from the ER lumen to the cytosol, independently of ER-associated degradation (ERAD), through a mechanism termed ER-to-cytosol signaling (ERCYS), which confers a selective advantage to tumor cells *(6)*. In parallel, oxidative stress, resulting from the generation of ROS and contributing to cellular damage, inflammation and impaired redox homeostasis, is an important determinant of cancer treatment efficacy *(4)*. Cellular adaptation to oxidative stress can protect cancer cells against anticancer drugs, suggesting an association between oxidative stress and drug resistance *(4)*. This adaptation involves the nuclear factor (erythroid 2)-like (NRF) transcription factors, such as NRF2/NFE2L2, which drive the expression of numerous genes involved in redox homeostasis *(7)*. Finally, proteome plasticity has emerged as a further mechanism enabling cells to cope with oncogenic signaling and microenvironmental stressors such as drug exposure. mRNA translation reshaping enables cells to rapidly remodel their proteome by modulating specific signaling pathways, thereby shaping cancer phenotypes *(5, 8)*. Selective translation reprogramming has been shown to enhance cancer cell survival and promote treatment resistance, primarily through the translation initiation complex eIF4F, which drives the translation of mRNAs associated with proliferation and stress adaptation *(9)*. Tight control of mRNA translation thus appears critical for counteracting cellular stresses and promoting cell survival in response to chemotherapy.

Molecular chaperones play a key role in the regulation of protein homeostasis and represent potential targets for overcoming chemoresistance *(10, 11)*. We previously identified the FKBP7 chaperone as overexpressed in taxane-resistant PCa cells and in prostate tumors, where its expression positively correlates with recurrence in patients receiving docetaxel *(12)*. FKBP7 is required for the growth of chemoresistant cell lines and tumors in mice *(12)*. This protein (UniProt Q9Y680 for the human protein) belongs to the FKBP family which comprises broadly expressed peptidyl-prolyl isomerases (PPIases) that accelerate the *cis-trans* isomerization of proline peptide bonds, a process involved in protein folding. The FKBP family includes the archetypical FKBP12 which binds the immunosuppressive drugs FK506 and rapamycin to inhibit respectively calcineurin and mTOR, through the formation of ternary complexes *(13, 14)*. FKBP7 belongs to the subgroup of ER-resident FKBPs characterized by an N-terminal ER-targeting sequence and a C-terminal ER-retention sequence. Like the other luminal ER-resident FKBPs, FKBP9, FKBP10 and FKBP14, FKBP7 contains two C-terminal EF-hands motifs with a helix-loop-helix topology that bind calcium *(14)*. The emerging role of ER-resident FKBPs in cancer progression has recently been highlighted by an increasing number of studies. We and others have reported overexpression of ER-resident FKBPs associated with poor prognosis and tumor recurrence in PCa *(12, 15)*, lung cancer *(16)*, glioblastoma *(17)*, or in melanoma *(18, 19)*. To date, FKBP7 remains poorly characterized both biochemically and structurally. Its isomerase catalytic activity has been demonstrated *in vitro* using synthetic substrates *(20, 21)*, and in murine cell lines, FKBP7 has been shown to modulate the ATPase activity of the ER-chaperone GRP78/BiP/HSPA5 through a calcium-dependent mechanism *(20, 22, 23)*. In our previous study *(12)*, we showed that FKBP7 interacts with the translation initiation complex eIF4F, suggesting a role for FKBP7 in translational control to promote the survival of resistant cells. The proteome remodeling potentially triggered by the FKBP7-eIF4F interaction could therefore be relevant to taxane-based chemoresistance in PCa.

Here, we sought to refine our understanding of how FKBP7 contributes to stress adaptation and taxane resistance in prostate cancer, with a particular focus on translation regulation. Specifically, we examined how FKBP7 integrates ER and oxidative stress signals in response to chemotherapy, characterized the structure of the FKBP7 PPIase domain and its regions of interaction with eIF4G1 within the translation machinery, and analyzed how FKBP7 silencing alters translation programs. Our results reveal an FKBP7-mediated adaptation mechanism involving nuclear factor (erythroid 2)-like (NRF) transcription factors that drive the oxidative stress response and may underlie chemoresistance.

## RESULTS

### N-glycosylated FKBP7 is recovered in the cytosol independently of proteasomal degradation

FKBP7 has been historically reported to localize in the lumen of the ER *(24)*. However, our previous work *(12)* showed that FKBP7 interacts with the eIF4G subunit of the eIF4F translation initiation complex. eIF4G1 is not itself an ER-resident protein and has not been reported to localize to the ER lumen; its interaction with FKBP7 therefore necessarily occurs outside the ER suggesting that FKBP7 can be exported from the ER to the cytosol. Such a mechanism could enable resistant cells to favor interaction between FKBP7 and eIF4G in the cytosol under specific conditions. To address this hypothesis, we carried out subcellular fractionation separating cytosolic proteins (Cyt.) from those contained in membrane organelles (Memb.), including ER proteins, using parental (S) and docetaxel-resistant (R) PCa cell lines. Due to the low expression of endogenous FKBP7, detection of cytosolic FKBP7 required prior protein precipitation of the fraction for enrichment. These results show that FKBP7 is present in the cytosol, both in sensitive and resistant cells. Although FKBP7 levels were much lower in the cytosolic fraction than in the ER, cytosolic FKBP7 was higher upon docetaxel treatment in both parental cell lines (S) though this increase did not reach statistical significance. In addition, cytosolic FKBP7 was significantly higher in resistant (R) than in parental IGR-CaP1 cells (S), with a similar trend observed in 22Rv1 cells (Fig. 1A), suggesting an association between cytosolic FKBP7 levels and docetaxel resistance.

**Figure 1:**
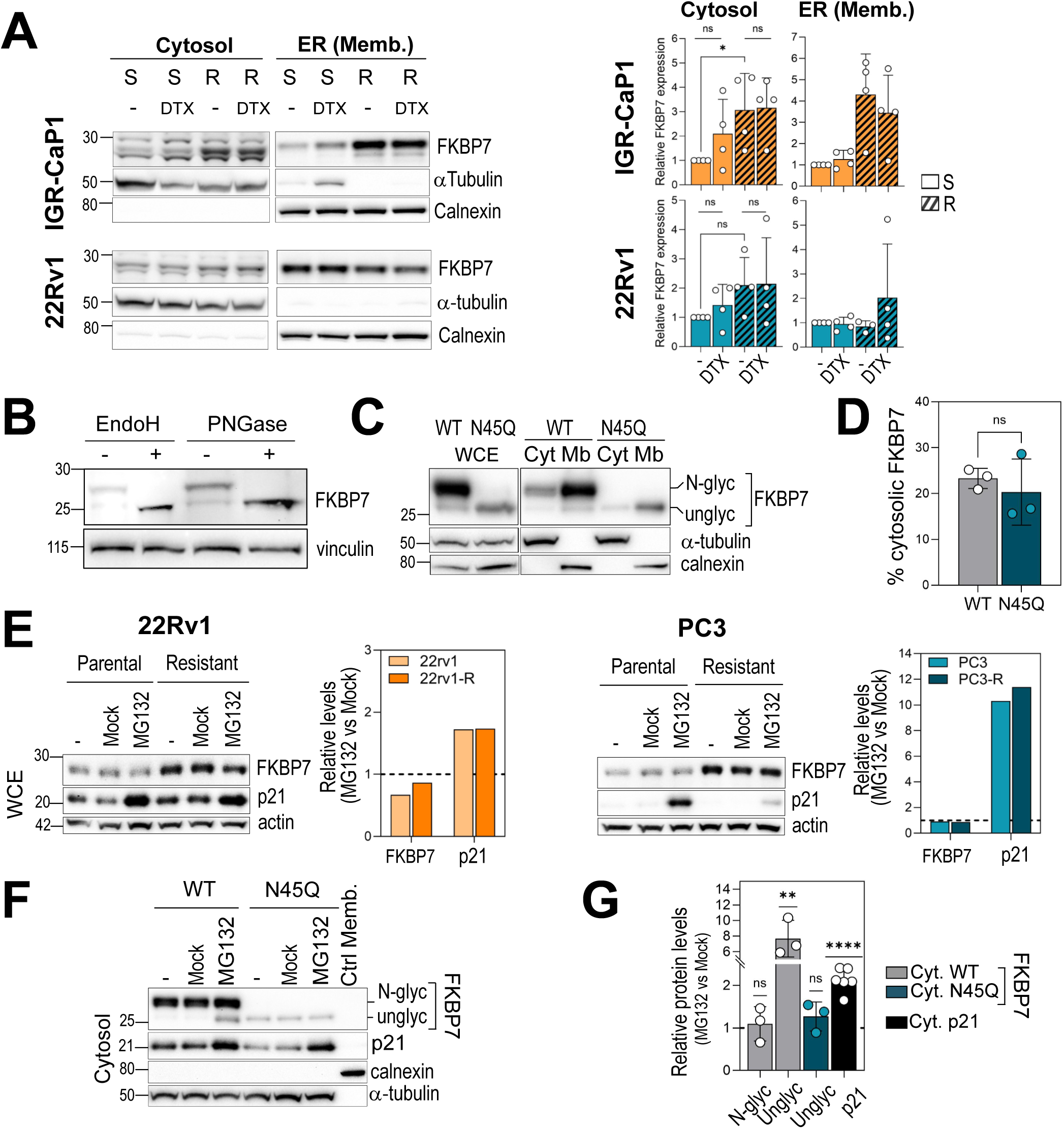
FKBP7 is refluxed into the cytosol independently of proteasome degradation. **A.** Immunoblot showing FKBP7 levels in cytosol (Cyt.) and membrane organelles (Memb.) in parental (S) or docetaxel (DTX)-resistant (R) IGR-CaP1 and 22Rv1 cell lines (R: hatched bars). Parental cells were left untreated or treated for 48-72 h with 5 nM DTX (22Rv1) or 10 nM DTX (IGR-CaP1), while DTX-resistant cells were left untreated or treated for 48-72 h with 125 nM DTX. Alpha-tubulin and calnexin are the fractionation controls for cytosol and membrane organelles respectively. Right: protein levels quantified using Image Lab software, expressed relative to the corresponding loading control and then relative to untreated parental cells, from four independent experiments. Data are mean ± SD. Statistical significance determined by ratio paired t-test: cytosolic fractions for R versus S comparisons (P=0.0358 for IGR-CaP1 cells; P=0.0736 for 22Rv1 cells); the effect of DTX effect did not reach statistical significance. **B.** Immunoblot showing the FKBP7 N-glycosylation status in LNCaP cells (whole cell extracts) after treatment or not with Endoglycosidase H (Endo H) or N-Glycosidase F (PNGase F). Vinculin is the loading control. **C**. Immunoblot showing identification of the FKBP7 N-glycosylation site after transfection of an expression vector encoding wild-type FKBP7 (WT) or the N45Q point mutant in 22Rv1 cells. Bands correspond to the N-glycosylated (N-Glyc) and non-glycosylated (Unglyc) forms. Whole cell extracts (WCE, left panel) and cytosolic (Cyt.) and membrane organelles (Memb.) fractions (right panel) are shown. Alpha-tubulin and calnexin are the fractionation controls for cytosol and membrane organelles respectively. **D**. Quantification of the cytosolic FKBP7 protein proportion (based on C) relative to its total amount (cytosol + membrane), from three independent experiments. Data are mean ± SD. Paired Student’s t-test; ns: non-significant (P=0.4555). **E**. Immunoblot showing total FKBP7 levels after treatment with MG132 (10 µM, 6h) or control in 22Rv1 and PC3 cells. p21 is the positive control for proteasome inhibition, actin is the loading control. Right: FKBP7 quantified relative to actin and expressed relative to the control (DMSO, Mock). **F.** Immunoblot showing cytosolic FKBP7 levels after treatment with MG132 (10 µM, 6h) or control in the 22Rv1 cells transfected with wild-type FKBP7 or N45Q mutant. p21 is the positive control for proteasome inhibition, alpha-tubulin and calnexin are the subcellular fractionation controls. **G.** Quantification from three independent experiments (six for p21) of signals normalized to the respective loading control, expressed relative to the untreated control (DMSO, Mock). Data are mean ± SD. Statistical significance was determined by one sample ratio t test.

Mouse FKBP7 has been shown to be N-glycosylated in the ER *(24)*. Protein N-glycosylation is a post-translational modification of proteins that initiates in the ER and can be modified in the Golgi apparatus. It favors protection of newly synthesized proteins from degradation *(25)*. To determine whether the cytosolic pool of human FKBP7 had passed through the ER before export to the cytosol, or is instead directly expressed in the cytosol in a non-glycosylated form, we assessed the glycosylation status of FKBP7 using Endoglycosidase H (EndoH) and N-Glycosidase F (PNGase) in LNCaP (Fig. 1B) and IGR-CaP1 cells (Fig. S1A). In both cell lines and with both enzymes, the upper band of the doublet shifted in size, confirming N-glycosylation of endogenous FKBP7, the non-glycosylated form migrating at 25 kDa. A single N-glycosylation site has been predicted at Asn45 of human FKBP7 using the NetNGlyc 1.0 server. A N45Q mutant lacking this putative Asn-glycosylation site was constructed and analyzed by whole cell extraction and cell fractionation. Unlike FKBP7-WT, the N45Q mutant showed only a single band at 25kDa in 22Rv1 (Fig. 1C) and PC3 cells (Fig. *S*1B), demonstrating that FKBP7-WT is N-glycosylated at the position N45. Subcellular fractionation of 22Rv1 expressing FKBP7-WT or FKBP7-N45Q further showed that the predominant cytosolic form of FKBP7-WT was N-glycosylated at position N45, as observed for ER-resident FKBP7 (Fig. 1C). Since glycosylation occurs in the ER, these results indicate that cytosolic FKBP7 originates from the ER rather than from direct cytosolic synthesis. They further indicate that N-glycosylation is not required for FKBP7 retro-translocation to the cytosol, as similar cytosolic fractions relative to total FKBP7 were observed for WT and N45Q mutant (Fig. 1D).

In case of protein misfolding or abnormal overproduction, ER-resident proteins can be retro-translocated to the cytosol to be degraded by the ERAD process *(26)*. We therefore evaluated if FKBP7 might be exported to the cytosol for degradation. Inhibition of the proteasome with MG132 in 22Rv1 and PC3 cells (Fig. 1E) showed that the total FKBP7 levels did not increase under MG132 treatment, in contrast to p21, a known proteasome target *(27)*, indicating that the bulk of FKBP7 pool is not targeted by the proteasome. However, to rule out possibility that changes in cytosolic FKBP7 levels upon MG132 treatment were masked by its low abundance in this compartment, cytosolic FKBP7 was analyzed more specifically under MG132 treatment. The results (Fig. 1F-G) showed that the non-glycosylated cytosolic form (WT Unglyc) accumulated under MG132 treatment, while the glycosylated form (WT N-glyc) remained unaffected. This indicates that the non-glycosylated cytosolic form of FKBP7 is degraded by the proteasome, whereas its cytosolic N-glycosylated counterpart, which is predominant, escapes this degradation. Thus, N-glycosylation appears to protect cytosolic FKBP7 from proteasomal degradation. We note that the unglycosylated N45Q mutant did not accumulate significantly upon MG132 treatment, possibly because it never engages the N-glycan-dependent ER quality control machinery *(25)* that normally acts on wild-type FKBP7.

Taken together, these results demonstrate that FKBP7 exhibits an atypical cytosolic localization, which is enhanced in docetaxel-resistant cells. The cytosolic form of FKBP7 is N-glycosylated prior its export from the ER and largely escapes proteasomal degradation. This suggests that FKBP7 has an unrecognized activity in the cytosol that might be linked to chemoresistance.

### The FKBP7 reflux into the cytosol is enhanced by chemotherapies

Since the cytosolic FKBP7 level is higher in docetaxel-resistant cells than in parental cells, we sought to determine to what extent this increase could be induced by docetaxel treatment in tumor cells. In PCa cells, the total FKBP7 level was previously shown to increase in response to low doses of docetaxel and cabazitaxel *(12)*. Here, cells were treated with several classes of chemotherapies, including the alkylating agent carboplatin (CPT), antimetabolites (5-FU and methotrexate (MTX)), and the topoisomerase inhibitor etoposide (ETO) at doses equivalent to the previously established IC₅₀ values (Fig. S2). These chemotherapies also induced an increase in total FKBP7 levels in both cell lines tested (Fig. 2A), suggesting that this response is not specific to taxanes but rather reflects a broader response to chemotherapy.

**Figure 2:**
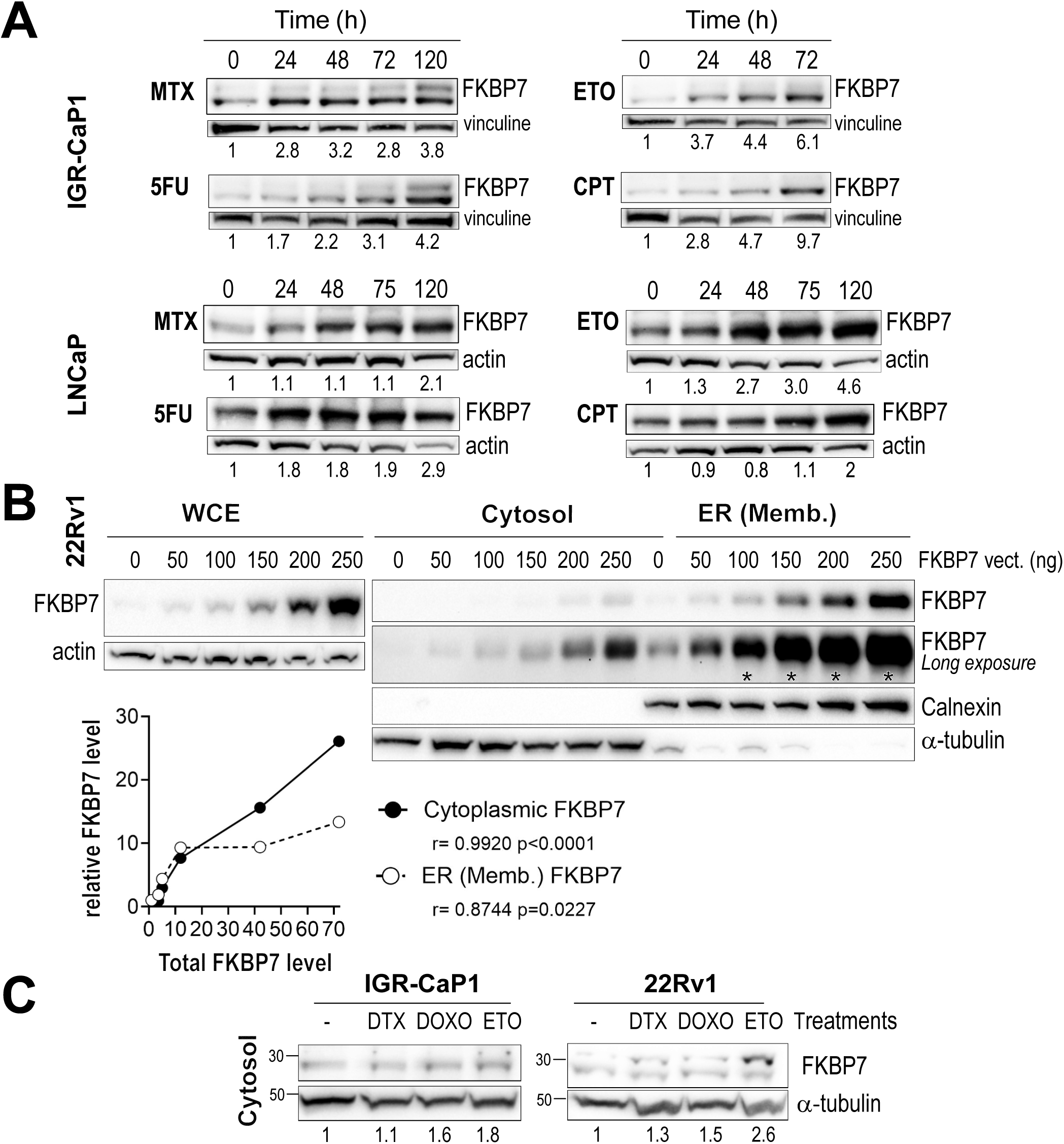
Cytosolic FKBP7 levels are enhanced by several chemotherapies. **A**. Immunoblots showing total FKBP7 levels in response to various chemotherapies in the IGR-CaP1 cells (methotrexate, MTX), 40 nM; 5-FU, 7.5 µM; etoposide (ETO), 6 µM; carboplatin (CPT), 34.5 µM) and LNCaP cells (MTX, 47 nM; 5FU, 1.1 µM; ETO, 2.4 µM; CPT, 50 µM), corresponding to the IC_50_ determined for each drug in the corresponding cell line (Fig. S2). Actin and Vinculin are the loading controls. Protein levels were quantified using Image Lab software; FKBP7 is expressed relative to the corresponding loading control and then relative to untreated cells (values shown below the blots). **B**. Immunoblot showing FKBP7 levels in 22Rv1 cells transfected for 48 h with increasing amounts of a wild-type FKBP7 expression vector, in whole cell extracts (WCE, left) or in cytosolic and the membrane organelle (Memb.) fractions after cell fractionation (right). Prolonged exposure was used to visualize cytosolic signals (*, saturated signals). Alpha-tubulin and calnexin are the subcellular fractionation controls; actin is the loading control for WCE. FKBP7 is quantified relative to the corresponding loading control. Below, linear regression between total FKBP7 (WCE) and cytosolic FKBP7/ER with Pearson’s r. **C.** Immunoblot showing cytosolic FKBP7 levels in cells treated or not for 48 h with docetaxel (DTX; 10 nM in IGR-CaP1, 5 nM in 22Rv1 cells), doxorubicin (DOXO; 100 nM in IGR-CaP1, 5 nM in 22Rv1), or etoposide (ETO, 2.5 µM). Alpha-tubulin is the fractionation control for cytosolic fractions. Protein levels, quantified using Image Lab software and expressed relative to the loading control and then to untreated cells, are shown below.

To determine whether the level of cytosolic FKBP7 reflux correlates with total FKBP7 levels, FKBP7 was assessed in both the cytosolic and membrane fractions of 22Rv1 cells transfected with increasing amounts of FKBP7. Cytosolic FKBP7 levels increased in a dose-dependent manner that fully correlated with total FKBP7 levels in whole-cell extracts (Fig. 2B), showing that FKBP7 reflux into the cytosol is strongly dependent on its overall expression level. Consistently, higher levels of cytosolic FKBP7 were observed to different extents in both IGR-CaP1 and 22Rv1 cells treated with the chemotherapeutic drugs doxorubicin (DOXO), etoposide (ETO), or docetaxel (DTX) (Fig. 2C). In contrast, enzalutamide, a second-generation androgen receptor-targeted therapy, had no effect on total FKBP7 levels in enzalutamide-sensitive or -resistant LNCaP-derived cells (Fig. S3), indicating a specificity of response to chemotherapy. Taken together, these results indicate that chemotherapy promotes FKBP7 reflux into the cytosol, primarily by increasing its global expression, rather than by inducing of specific retrotranslocation mechanism.

### Enhancement of FKBP7 cytosolic reflux upon chemotherapy is associated with an increased oxidative stress

Cytotoxic chemotherapies cause cellular stress, which can result in cell death. The ER plays a key role in this process, as ER stress and activation of the UPR pathway contribute to chemotherapy resistance *(28)*. These treatments also increase ROS production, resulting in oxidative stress *(4)*. Together, ER and oxidative stresses present major challenges that cancer cells must overcome to survive and resist therapy. We therefore focused on these two types of stress to identify the chemotherapy-induced cellular processes associated with increased cytosolic FKBP7 retrotranslocation. Tunicamycin (TN) and thapsigargin (TG), described as strong inducers of ER stress, and brefeldin A (BFA), an inhibitor of ER-Golgi protein trafficking, were used to induce ER stress. Dose-dependent treatment with these three agents indeed induced ER stress activation in 22Rv1 cells ectopically overexpressing FKBP7, as evidenced by increased expression or phosphorylation of UPR pathway markers from the IRE1α, PERK and ATF6 branches, although a lower effect was observed for tunicamycin (Fig. 3A). In contrast, docetaxel and cabazitaxel did not activate any of the three UPR branches (IRE1α, PERK, ATF6) relative to control condition (Fig. 3B). Only etoposide activated the IRE1α branch without inducing PERK or ATF6 markers (Fig. 3B), indicating that UPR activation is not a shared feature of the chemotherapies tested, although all of them have been shown to increase of FKBP7 levels (*(12)*, Fig.2A). To rule out the possibility that ER stress activation contributes to taxane resistance, we further evaluated the effect of the GRP78/BiP inhibitor HA15, previously shown to inhibit cancer cell survival and overcome drug resistance *(29, 30)*, on viability of resistant cells. High doses of HA15 reduced viability of 22Rv1-R cells (Fig. 3C), whereas IGR-CaP1-R cells were largely unaffected (Fig. 3C). While the ER stress response differed between cell lines, ER stress does not appear to play a consistent role in taxane chemoresistance in our models.

**Figure 3:**
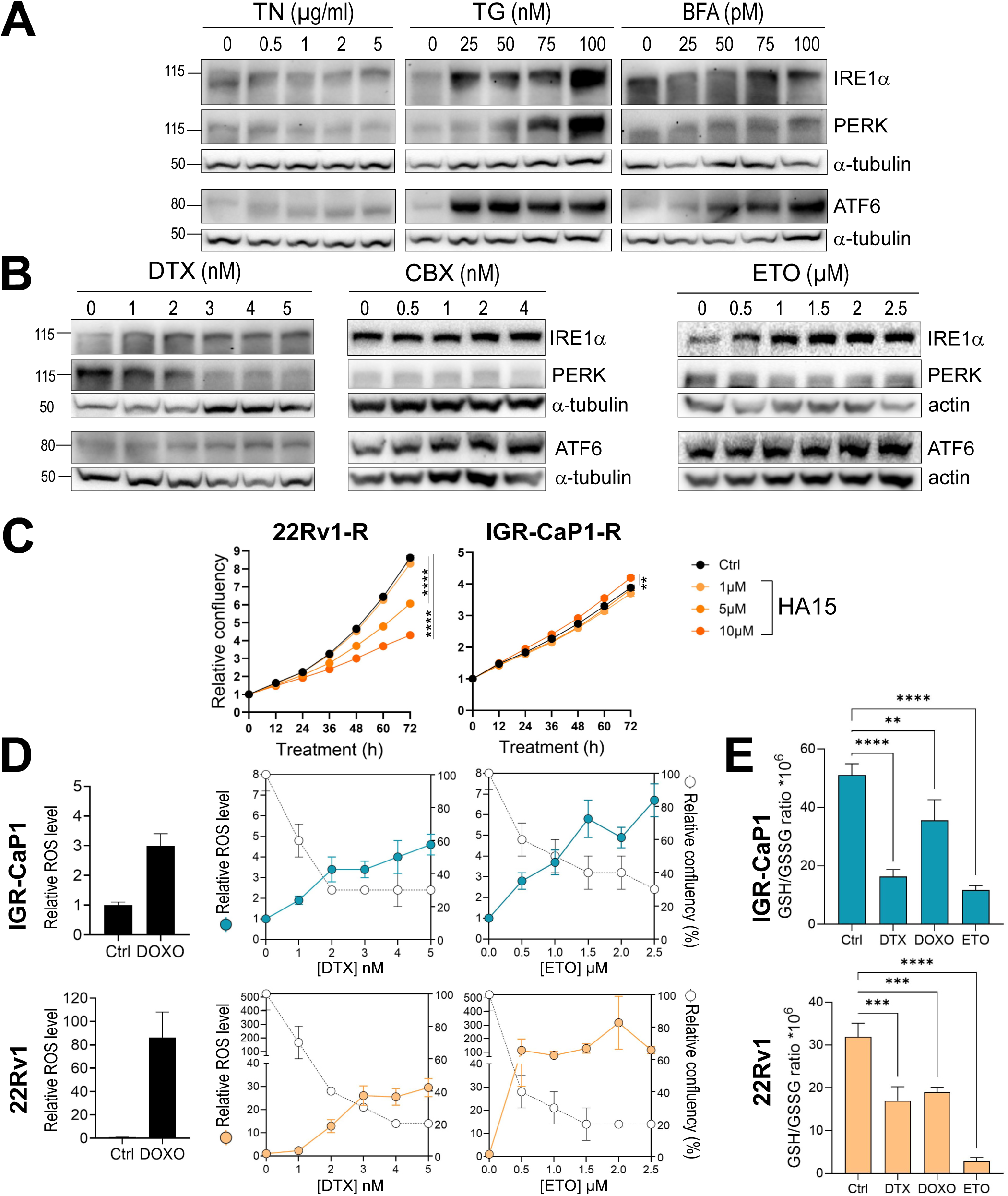
Reflux of FKBP7 into the cytosol is potentiated in response to oxidative stress. **A.** Immunoblot assessment of ER stress markers of the IRE1α, PERK and ATF6 branches after 16h treatment with increasing concentrations of tunicamycin (TN), thapsigargin (TG), or brefeldin A (BFA) in 22Rv1 cells transfected with 250ng of FKBP7-expressing vector. Alpha-tubulin is the loading control. **B.** Immunoblot showing ER stress markers as in A, in cells treated with increasing concentration of docetaxel (DTX), cabazitaxel (CBX) or etoposide (ETO) for 48 h. **C.** Cell proliferation of docetaxel-resistant IGR-CaP1 and 22Rv1 cells treated with increasing doses of ER stress inhibitor HA15 (orange) or vehicle control (black). Data are mean ± SEM. ** P < 0.01; **** P < 0.0001, 2-way ANOVA with Dunnett’s post-tests (at 72h). **D.** ROS levels (H_2_DCFDA, colored circle) and cell confluency (empty circle) were determined simultaneously in IGR-CaP1 and 22Rv1 cells treated with increasing doses of DTX or ETO for 48 h, or with 100 nM of doxorubicin (DOXO) for 48 h as positive control. ROS levels are expressed as Integrated Intensity Average per object normalized to confluence, and normalized to the untreated (Mock) condition. Data (representative of two independent experiments) are mean ± SEM. **E.** GSH/GSSG ratio after 48 h treatment with DTX (10nM in IGR-CaP1, 5 nM in 22Rv1), DOXO (100 nM in IGR-CaP1, 10 nM in 22Rv1) or ETO (5 µM in IGR-CaP1, 1 µM in 22Rv1). Data (representative of two independent experiments) are mean ± SD. ** P <0.01; *** P < 0.001; **** P < 0.0001, one-way ANOVA with Dunnett’s post-hoc test.

Cytotoxic chemotherapies such as platinum-based compounds and anthracyclines are known to generate high levels of ROS, leading to apoptosis *(31)*. In prostate cancer, oxidative stress has been associated with the response to therapy without a thorough understanding of the molecular biology involved *(32)*. Here, doxorubicin induced high levels of ROS in both the IGR-CaP1 and 22Rv1 cell lines, albeit to different extents (Fig. 3D). Increasing doses of docetaxel and etoposide also induced ROS production in our models within the same order of magnitude, with maximum ROS levels correlating with cytotoxicity (Fig. 3D). A reduction in the GSH/GSSG ratio in the presence of these chemotherapies further supports their ability to trigger oxidative stress (Fig. 3E). Conversely, no increase in ROS production was observed in IGR-CaP1 docetaxel-resistant cells treated with doxorubicin, docetaxel or etoposide (Fig. S4A). In 22Rv1 resistant cells, docetaxel similarly failed to increase ROS production, whereas etoposide and doxorubicin induced a modest increase, markedly lower than in parental cells (Fig. 3D and Fig. S4A). Consistently, no significant decrease in the GSH/GSSG ratio was observed upon chemotherapy treatment in either resistant cell line (Fig. S4B).

Overall, our results showed that docetaxel and a panel of chemotherapies increase total FKBP7 expression (Fig. 2A, *(12)*), that docetaxel, etoposide and doxorubicin increase cytosolic FKBP7 levels (Fig. 2C), and that these treatments also increase ROS production and decrease GSH/GSSG ratio (Fig. 3D-E). Altogether, these results indicate that resistant cells have developed the ability to counteract chemotherapy-induced oxidative stress by suppressing ROS production, together with an increase in FKBP7 cytosolic levels following chemotherapy exposure.

### FKBP7 interacts with the central region and the C-terminal HEAT3 structured domain of eIF4G1

Our previous work showed that FKBP7 interacts with the eIF4G scaffolding subunit of the eIF4F translation initiation complex *(12)*. The eIF4G1 isoform is one of three existing eIF4G isoforms and has been shown to be the most abundant in tumors *(33)*; it also plays an essential role in PCa *(34)*. eIF4G1 is a protein composed of 1600 amino acids with a flexible and unstructured N-terminal extremity, and central and C-terminal moieties that contain structured HEAT domains (HEAT1, HEAT2, HEAT3) composed of repeating helix-turn-helix motifs that govern interactions with eIF4A, eIF3 and MNK1/2 *(35, 36)* (Fig. 4A).

**Figure 4:**
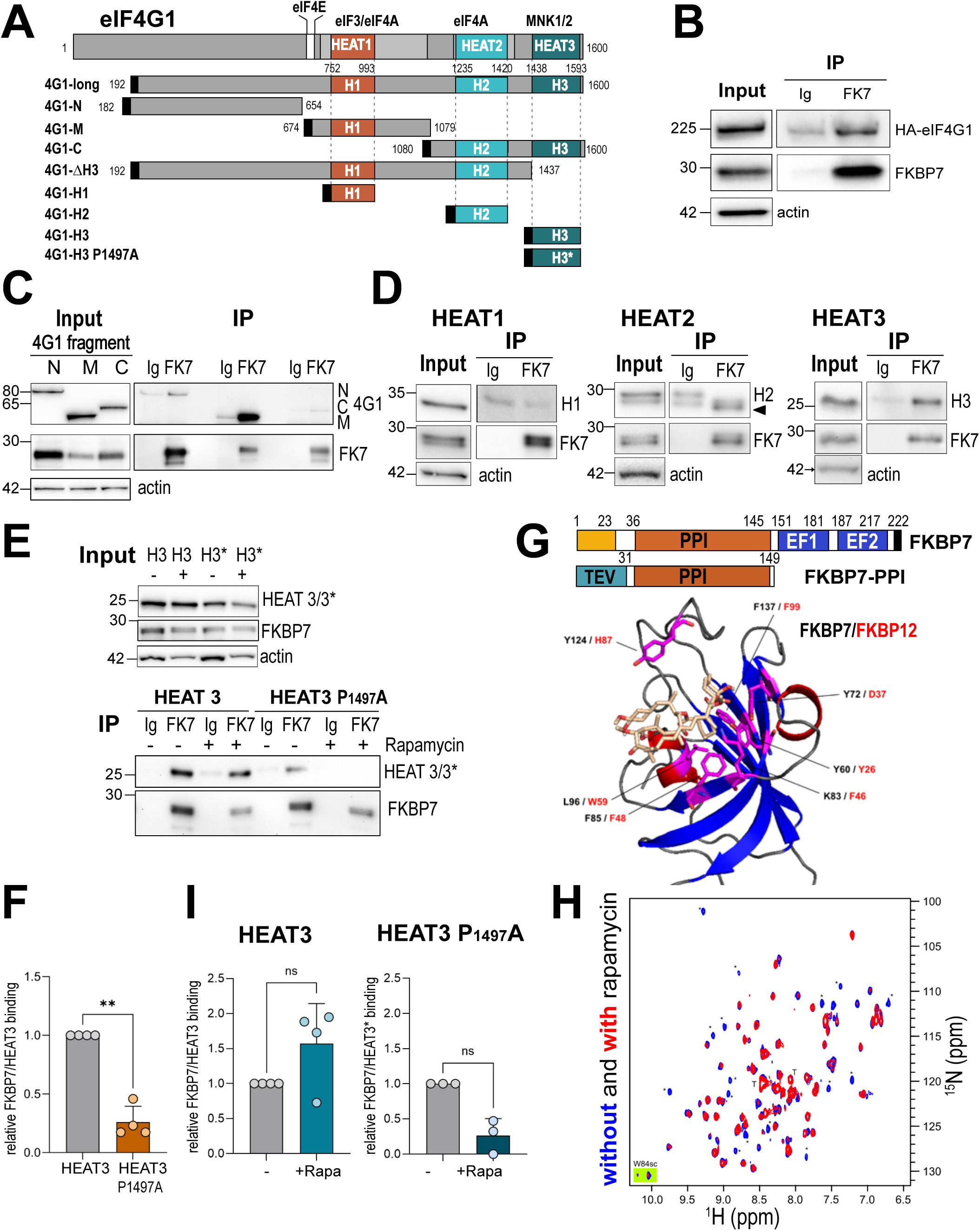
FKBP7 binds to rapamycin and interacts with the structured domains of eIF4GI. **A.** Schematic representation of eIF4G1 domains and the series of HA-tagged eIF4G1 constructs (black box). The three structured HEAT domains are highlighted in color. **B.** Immunoblot showing FKBP7 and co-immunoprecipitated eIF4G1 (4G1-long), revealed with an anti-HA antibody. FKBP7 was immunoprecipitated using an anti-FKBP7 antibody (or IgG as control) in PC3 cells co-transfected with FKBP7-WT and HA-eIF4G1. Input controls (10%) are shown. **C.** FKBP7 was immunoprecipitated as in B. Immunoblot showing FKBP7 and coimmunoprecipitation of the N-, M-, and C-terminal regions of eIF4G1, revealed with an anti-HA antibody. Input controls (10%) are shown. **D.** Immunoblot showing FKBP7 and coimmunoprecipitation of the isolated HEAT domains of eIF4G1, revealed with the anti-HA antibody. For the HEAT2, the arrow indicates the residual FKBP7 signal. Input controls (10%) are shown. **E.** Immunoblot showing FKBP7 and coimmunoprecipitated HEAT3 (H3) domains of eIF4G1, either wild-type or bearing the Pro1497Ala mutation (*), with or without pre-incubation of cells with 500 nM rapamycin for 30 min. Immunoblots were revealed with anti-HA and anti-FKBP7 antibodies. Input controls (10%) are shown. **F.** Quantification FKBP7/HEAT3 interaction, wild-type versus P1497A (based on E) from four independent experiments. Data are mean ± SD. Ratio paired Student’s t-test (P=0.0082)**. G.** Top, schematic representation of FKBP7 domains and of the PPIase construct bearing a TEV signal for recombinant protein production. Bottom, structural prediction of the FKBP7 PPIase domain generated with Phyre2 based on the FKBP14 crystal structure (PDB 4MSP), superimposed with the experimental structure of the FKBP12-rapamycin complex (PDB 1FKB), showing the predicted rapamycin-binding site. FKBP7 numbering in black, FKBP12 numbering in red. **H.** ^15^N SOFAST-HMQC NMR spectrum of ^15^N-labeled FKBP7 PPI domain alone (blue) and in the presence of a 1:1 molar ratio of rapamycin (red). Each cross-peak corresponds to one H-N chemical bond, approximately one amino acid residue. A few peaks, such as the Try84 side chain (green background), are doubled, suggesting multiple FKBP7 conformations in solution. Most, but not all, cross-peaks showed reduced intensity upon rapamycin addition, and about 15 cross-peaks (starred) disappeared completely. Peak intensity loss is quantified in Fig S6B. **I.** Quantification of the effect of rapamycin on the FKBP7/HEAT3 interaction, wild-type versus P1497A (based on E), from 3-4 independent experiments. Data are mean ± SD. Ratio paired Student’s t-test (ns, non significant).

We aimed to identify the interaction site(s) between FKBP7 and eIF4G1 to determine which of eIF4G1’s functions and/or interactions might be affected by its association with FKBP7. To achieve this, we performed immunoprecipitation experiments using various eIF4G1 constructs, presented in Fig. 4A, after expression in PC3 cells. Firstly, we demonstrated that FKBP7 interacts with the long eIF4G1 isoform (Fig. 4B). Secondly, we analyzed the interaction between FKBP7 and different eIF4G1 domains by using large deletion mutants as defined in Fig. 4A. Our results revealed a robust interaction with the central domain of eIF4G1 (eIF4G1-M), which harbors interaction sites for eIF4A and eIF3 and contains the HEAT1 domain (Fig. 4C). To a lesser extent, FKBP7 also appeared to interact with the 4G1-N and 4G1-C constructs of eIF4G1 (Fig. 4C). We next investigated the interaction between FKBP7 and the structured HEAT domains of eIF4G1 independently. Our results revealed a strong interaction between FKBP7 and the isolated C-terminal HEAT3 domain (Fig. 4D), while we were unable to confirm any interaction with the isolated HEAT1 and HEAT2 domains at this stage, despite HEAT1 being located within the eIF4G1-M construct (Fig. 4D). However, deletion of the HEAT3 domain from eIF4G1 did not result in the loss of the FKBP7/eIF4G1 interaction (Fig. S5A). Taken together, these results demonstrate that multiple regions of eIF4G1 are required for interaction with FKBP7. Consistently, since FKBP7 strongly interacted with the central domain (eIF4G1-M) but not with a construct restricted to the HEAT1 domain (Fig. 4D), one or more additional docking sites likely exist in fragments 674-752 and 993-1079, which flank the HEAT1 domain within eIF4G1-M. Finally, analysis of the published crystal structure of the eIF4G1 HEAT3 domain (PDB 1UG3) *(37)* revealed a proline at position 1497 (P1497) in the *cis* configuration instead of the more common *trans* configuration. Given the peptidyl-prolyl isomerase activity of FKBP7 *(20)*, we investigated whether this proline might be involved in the interaction between FKBP7 and the HEAT3 domain. Mutation of Pro1497 to alanine significantly reduced FKBP7 binding to HEAT3 (75% loss of interaction) (Fig. 4E-F), confirming its role in FKBP7/HEAT3 interaction. It is then possible that the presence of an alanine instead of a proline prevents FKBP7-mediated *cis-trans* isomerization at this position, with an impact on the HEAT3 interaction network or its folding, although other structural mechanisms unrelated to isomerization are also possible. Nevertheless, these data univocally show that Pro1497 has an important role in HEAT3/FKBP7 interaction. In summary, this detailed study of the FKBP7/eIF4G1 interaction indicates that FKBP7 interacts with several regions of eIF4G1, including, but not restricted to the C-terminal HEAT3 domain, where isomerization of the proline peptide bond at Pro1497 by FKBP7 could stabilize the FKBP7-eIF4G1 interaction in this region.

The interaction between FKBP7 and eIF4G1 in the cytosol suggests that FKBP7 could influence eIF4F-dependent translation initiation, thereby potentially impacting the proteome of resistant cells and promoting survival. We therefore investigated whether some ligands could interfere with FKBP7 binding to eIF4G1 to block this process. As there are no available experimental structures or specific ligands for FKBP7, we first predicted its structure by homology using Phyre2 *(38)* and AlphaFold *(39)*. There is high sequence conservation between FKBP PPIase domains in general and between FKBP7 and FKBP14 in particular (with 52% identity). FKBP14 also shares the same domain organization as FKBP7 with two EF-hands. We then generated a robust, high-quality structural prediction of FKBP7 PPIase (Fig. 4G) based on the FKBP14 crystal structure (PDB 4MSP) *(40)*. Rapamycin and FK506 are well-known FKBP12 ligands that also interact with other FKBPs *(14)*. Figure 4G shows our predicted structure of the FKBP7 PPIase domain and the potential rapamycin binding site, derived by homology to the FKBP12-rapamycin experimental structure (PDB 1FKB).

We then experimentally evaluated the interaction of rapamycin, everolimus (a rapamycin analog), and FK506 with the FKBP7 catalytic domain as potential tools to disrupt the FKBP7/eIF4G1 interaction. To this end, the recombinant ^15^N-labeled catalytic domain of FKBP7 (^15^N-FKBP7-PPI) was produced in *E. coli* and purified by affinity chromatography (Fig. 4G). Our data first show that the ^1^H-^15^N correlation NMR spectrum of ^15^N-FKBP7-PPI (shown in blue in Fig. 4H) is characteristic of a well-folded protein.

Additionally, NMR titrations of ^15^N-FKBP7-PPI with rapamycin revealed a loss of specific signals in the ^1^H-^15^N correlation spectrum of the FKBP7 catalytic domain (Fig. 4H in red), indicating that FKBP7 binds to rapamycin. Of note, rapamycin was dissolved in DMSO, and control NMR experiments showed that DMSO alone did not impact FKBP7 structure and induced only minor, non-specific peak intensity changes in 2D ^1^H-^15^N NMR spectra (Fig. S6A,D) that could not account for the pattern of signal loss observed upon rapamycin or everolimus binding. Typically, the formation of a protein-ligand complex is associated with peak shifts in protein NMR spectra. In this case, however, the disappearance of specific cross-peaks upon rapamycin binding most likely suggests multiple interconverting conformations for rapamycin within the binding site. In the absence of peak assignment, the localization of rapamycin on FKBP7 PPIase was not further explored. Everolimus produced similar results to rapamycin (Fig. S6C,D), suggesting a comparable binding mode. In contrast, no NMR spectral modification was observed upon the addition of FK506, indicating an absence of intermolecular interaction with FKBP7 under the evaluated conditions (Fig. S6B,D).

We next evaluated the ability of rapamycin to alter the interaction between FKBP7 and the HEAT3 domain in cells. Treating cells with an excess of rapamycin did not significantly alter the interaction between FKBP7 and either the wild-type or the P1497A mutant HEAT3 domain (Fig. 4E,I), although the already weaker HEAT3-P1497A binding appeared more sensitive to rapamycin than wild-type binding. While the exact binding site of rapamycin and everolimus on FKBP7 was not firmly determined, both ligands most likely bind and block the pocket for proline-containing substrates, as classically observed with other FKBPs.

### FKBP7 interacts with the translational machinery and contributes to translation

The interaction between FKBP7 and eIF4G1 suggests that FKBP7 may be involved in translation initiation, particularly in the context of docetaxel resistance. To analyze the translational status of mRNAs, we performed polysome profiling experiments in parental and docetaxel-resistant IGR-CaP1 (Fig. 5A) and 22Rv1 (Fig. S7A) cell lines. FKBP7 co-sedimented with fractions containing 40S small ribosomal subunits, which also contained eIF4G1. Supporting this observation, FKBP7 co-immunoprecipitated with RPS6, a component of the 40S small ribosomal subunit (Fig. 5B). FKBP7 was also detected, to a lower extent, in polysome fractions, suggesting a potential involvement in translation elongation (Fig. 5A and Fig. S7A-B). To rule out that FKBP7 detection in polysome fractions reflects contamination by ER-derived material rather than bona fide association with cytosolic ribosomal complexes, polysome profiling was performed on cytosolic extracts obtained after selective permeabilization of the plasma membrane with digitonin to exclude ER-derived structures. FKBP7 was detected in fractions containing 40S ribosomal subunits and polysomes (Fig. S7B). The absence of calnexin and RPN1, a membrane-bound component of the rough ER, in all fractions excluded ER-derived contamination (Fig. S7B). EDTA treatment, which dissociates polysomes into ribosomal subunits, abolished the FKBP7 signal in the polysome fraction (Fig. S7B), consistent with FKBP7 sedimentation depending on intact ribosomal assembly rather than on nonspecific aggregates. These results are consistent with our previous mass spectrometry and SILAC data showing an association of FKBP7 with components of the eIF2, eIF4 and mTOR signaling pathways *(12)*, and support a model in which FKBP7 participates in translation pre-initiation and/or initiation, particularly in docetaxel-resistant PCa cells that overexpress FKBP7 in the cytosol.

**Figure 5:**
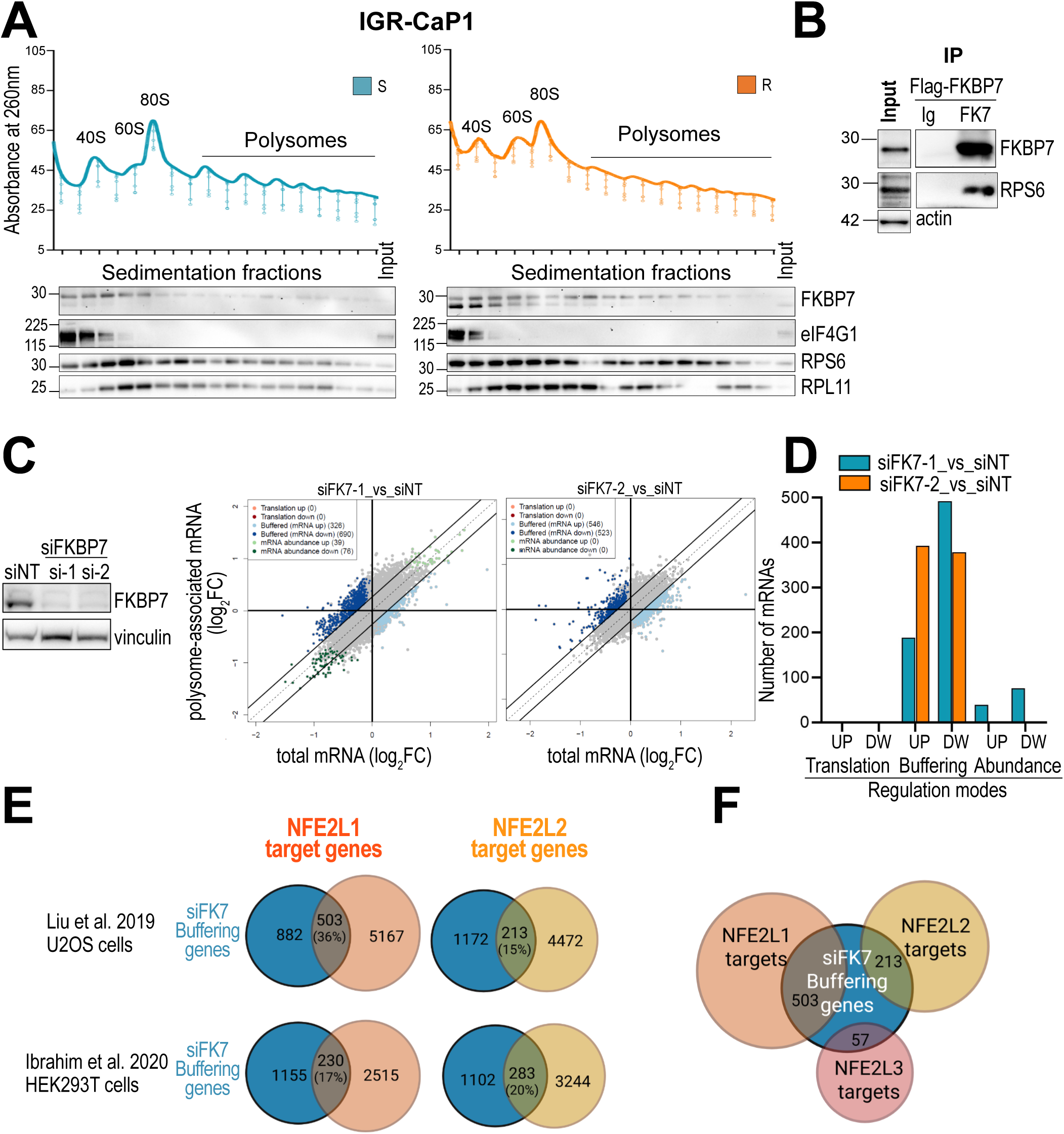
FKBP7 interacts with the ribosomal subunits and contributes to translation. **A.** Polysome profiles of parental (S, blue) and docetaxel-resistant (R, orange) IGR-CaP1 cells under basal conditions (representative of three independent experiments). Immunoblot showing FKBP7, eIF4G1, RPS6 (40S ribosomal subunit marker) and RPL11 (60S ribosomal subunit marker) in each fraction. Input corresponds to the cellular extract before loading onto sucrose gradient. **B.** FKBP7 was immunoprecipitated using an anti-FKBP7 antibody (or IgG as control) in PC3 cells transfected with a Flag-FKBP7-WT encoding vector. Immunoblot showing FKBP7 and co-immunoprecipitated RPS6. Input controls (10%) are shown. **C.** Left, immunoblot showing FKBP7 knockdown efficiency 48 h after transfection with two FKBP7-targeting siRNAs in docetaxel-resistant IGR-CaP1 cells, used for polysome profiling analysis. Vinculin, loading control. Right, scatter plots of polysome-associated versus total cytosolic mRNA log_2_ fold changes (siFK7-1 vs. siNT and siFK7-2 vs. siNT), colored according to their regulatory mode as derived from anota2seq analysis of RNA-seq data from polysome profiling in resistant IGR-CaP1 cells (obtained from three independent experiments). **D.** Number of translated (translation), transcribed (mRNA abundance), and buffered mRNAs (FDR < 0.05), based on analysis shown in C. **E.** Comparison of the 1,391-gene set identified as translationally buffered (siFK7 vs. siNT, Fig. 5D and Dataset S2) with independent signatures of NFE2L1 and NFE2L2 target genes (44, 45). **F**. Overlap between NFE2L1, NFE2L2 and NFE2L3 target genes from the Liu *et al.* signature (44) and the translationally buffered gene set.

To explore the impact of FKBP7 on translation regulation, we next examined the effect of FKBP7 silencing on the translatome of docetaxel-resistant cells by performing RNA sequencing following polysome profiling experiments in IGR-CaP1-R cells transiently silenced for FKBP7 using two different siRNAs (siFKBP7-1 and siFKBP7-2), or treated with a control siRNA (siNT) (Fig. 5C). The anota2seq algorithm, used here for differential gene expression analysis, classifies transcripts into three regulatory modes: abundance, where changes in polysome-associated mRNAs are proportional to changes in total mRNAs; translation, where changes in polysome-associated mRNAs are not explained by changes in total mRNAs; and translational buffering, where ribosome occupancy offsets the effects of transcriptional variation *(41)*. Comparing fold changes in total and polysome-associated mRNAs upon FKBP7-silencing, no transcripts were classified in the translation category, indicating that FKBP7 silencing did not induce changes in translation efficiency independently of total mRNA levels. Only a small number of transcripts were proportionally modified at the total and polysome-associated mRNA levels (39 mRNA Up and 76 mRNA Down, FDR <0.05, abundance category) for one of the two siRNAs (Fig. 5C-5D, Fig. S7C and Tables S2A-B). In contrast, we identified a large population of translationally buffered mRNAs (581 up and 871 down, FDR <0.05) corresponding to a set of 1,391 non-redundant annotated genes (Fig. 5C-D and Fig. S7B, Tables S2A-B and S3). Gene Set Enrichment Analysis (GSEA) of the complete translationally buffered gene set (2,085 genes) identified enrichment of metabolic pathways, including “cholesterol biosynthesis pathway” and “fatty acids and lipoproteins transport”, together with upregulation of the NRF2 pathway and downregulation of DNA repair pathways (Fig. S8A). Reactome Pathways Enrichment analysis confirmed the upregulation of metabolic pathways and the downregulation of DNA repair and cell cycle pathways (Fig. S8B).

The transcription factor NRF2/NFE2L2 and, more recently its homologue NFE2L1, have been reported as master regulators of oxidative stress and redox homeostasis maintenance *(7, 42, 43)*. NFE2L2 has also been shown to mediate adaptation to oxidative stress induced by chemotherapy or radiotherapy *(7)*. We therefore compared our buffering gene set (1,391 genes) with NFE2L1 and NFE2L2 target genes identified in human cells by ChIP-seq *(44)* or by transcriptomic and proteomic analyses *(45)*. Venn diagrams revealed a consistent overlap between our buffering gene set and NFE2L1 targets (36% and 17% of the buffering genes, respectively, in each study) and NFE2L2 targets (15% and 20% respectively) (Fig. 5E). Using the signature from Liu et al. *(44)*, the largest number of target genes was found for NFE2L1 (503 genes), compared with NFE2L2 (213 genes) and NFE2L3 (57 genes) (Fig. S9A). Considering anota2seq regulation categories, 56% of the buffering genes (Fig. 5F) and 44% of the genes modified at the transcriptional level (Fig. S9A) upon silencing of FKBP7 corresponded to target genes of NRF (NFE2L1/2/3) transcription factors. Taken together, these results demonstrated that FKBP7 silencing affects these transcription factor pathways in docetaxel-resistant cells, and more specifically the NFE2L1 pathway.

Regarding a direct regulation of NFE2L1/NFE2L2 by FKBP7, a direct effect on their transcription appears unlikely, as FKBP7 silencing in resistant cells did not significantly modify either NFE2L1 nor NFE2L2 transcription, as shown in our anota2seq analysis (Tables S2A-B), and by RT-qPCR (Fig. S9B). Our anota2seq analysis also did not reveal any direct alteration in the translation of NFE2L1 or NFE2L2 transcripts themselves upon FKBP7 silencing (Tables S2A-B). Nevertheless, the large overlap between the buffering gene set and NFE2L1/NFE2L2 target genes suggests that FKBP7 could affect NFE2L1 and/or NFE2L2 protein activity, stability, or translation. The absence of a detectable effect on NFE2L1/NFE2L2 translation in our dataset may also reflect an early, transient regulatory event no longer detectable after 48 hours of FKBP7 silencing.

### FKBP7 increases NFE2L1 protein levels

To evaluate the direct impact of FKBP7 on NFE2L1 and NFE2L2, we next examined the protein levels of these two transcription factors by immunoblot under conditions of FKBP7 overexpression or silencing. These transcription factors undergo a complex activation process in the cytosol before becoming transcriptionally active in the nucleus *(7, 46)*. The newly synthesized forms of NFE2L1 and NFE2L2 are glycosylated and migrate at ∼100kDa. In IGR-CaP1 and 22Rv1 cells, these forms were only detectable upon treatment with the proteasome inhibitor MG132 (Fig. 6A), owing to their short half-life *(47, 48)*. In both cell lines, NFE2L1 levels increased in resistant compared to parental cells in the presence of MG132, whereas glycosylated NFE2L2 was either undetectable or present at equal levels in sensitive and resistant cells (Fig. 6A).

**Figure 6:**
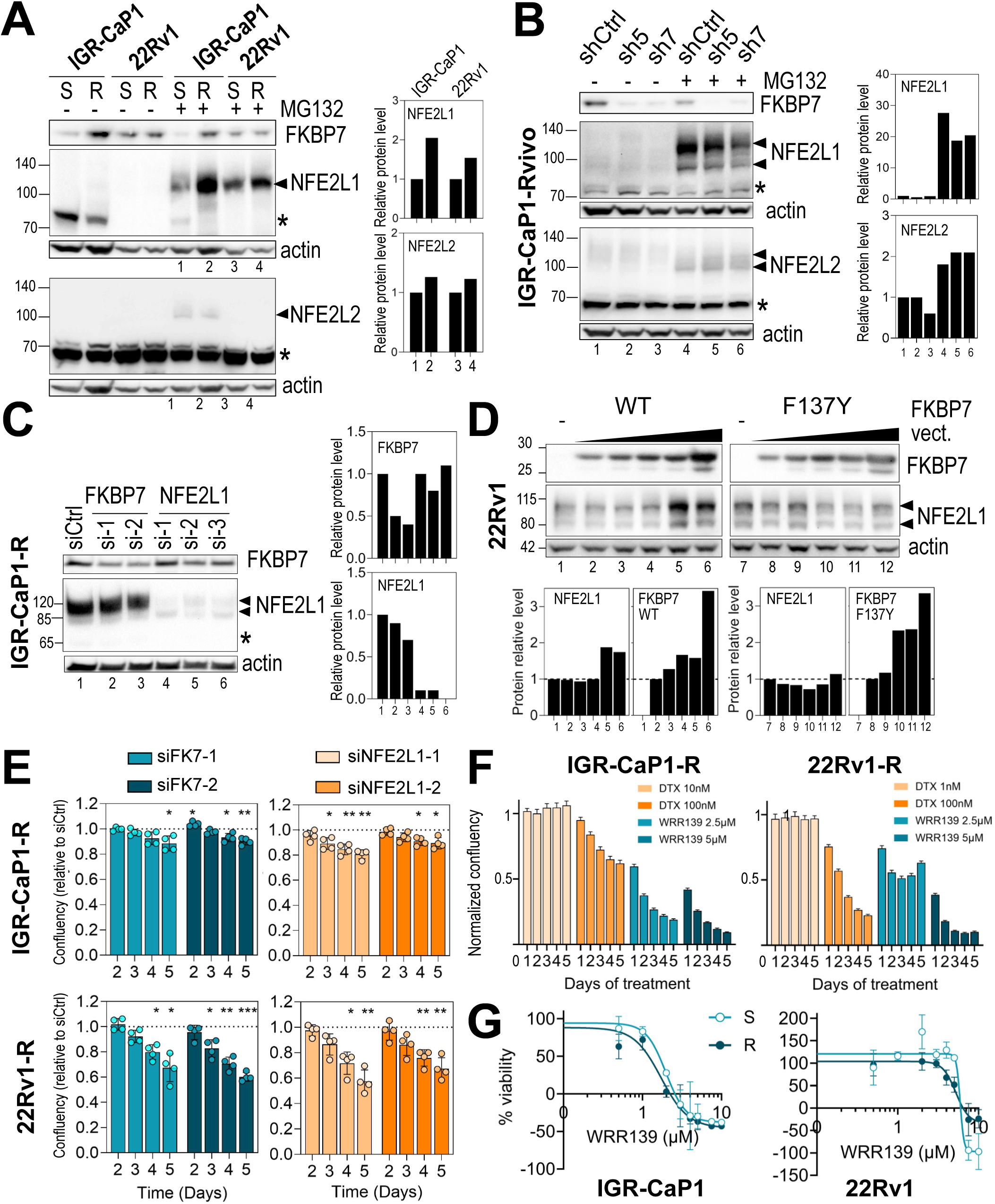
FKBP7 increases NFE2L1 protein levels. **A**. Immunoblot showing FKBP7, NFE2L1 and NFE2L2 levels in whole cell extracts from parental (S) or docetaxel-resistant (R) IGR-CaP1 and 22Rv1 cells, treated or not with 10 µM MG132 for 24 h, representative of three independent experiments. Actin is the loading control. Only the high-molecular-weight (85-150 kDa) forms of NFE2L1 and NFE2L2, corresponding to newly synthesized protein (arrows), are quantified; nuclear forms of NFE2L1 and NFE2L2 are indicated by stars. Right, NFE2L1 and NFE2L2 levels relative to actin and to parental cells, from the blot shown. **B**. Immunoblot as in A, in IGR-CaP1-Rvivo cells, a docetaxel-resistant IGR-CaP1 subline selected *in vivo*, depleted for FKBP7 (sh5 and sh7) or not (shCtrl). Protein level quantification as in A, from the blot shown. **C**. Immunoblot showing FKBP7 and NFE2L1 in whole cell extracts from resistant IGR-CaP1 cells transfected with siRNA targeting FKBP7 or NFE2L1 for 48 h, treated with 10 µM MG132 for the last 24 h; representative of four independent experiments. Right, quantification as in A, from the blot shown. **D**. Immunoblot showing FKBP7 and NFE2L1 after 48-72 h transfection with increasing amounts of wild-type or Phe137Tyr (F137Y) mutant FKBP7 expression vector in 22Rv1 cells, treated with 10 µM MG132 for the last 24 h; representative of two independent experiments. Below, quantification as in A, from the blot shown. **E**. Daily cell proliferation, determined by live imaging, after transfection with different siRNA targeting FKBP7 (blue) or NFE2L1 (orange) in two resistant cell lines. Data (four independent experiments) are mean ± SD, normalized to the siNT control. * P < 0.05; ** P < 0.01; *** P < 0.005, 2-way ANOVA with Dunnett’s post-tests. **F.** Daily cell proliferation, determined by live imaging, after treatment with docetaxel (1-100 nM, orange bars) or the NGLY1/NFE2L1 inhibitor WRR139 (2.5 and 5 µM, blue bars) in two docetaxel-resistant cell lines. Data (representative of three experiments) are mean ± SEM, normalized to the Mock control. **G.** Cell viability of parental and docetaxel-resistant IGR-CaP1 and 22Rv1 cells after treatment with increasing doses of WRR139 for 48 h. Data are mean ± SD. IC_50_ values, calculated using four-parameter logistic regression (GraphPad), were 1.72 µM (resistant) vs. 1.98 µM (parental) in IGR-CaP1 cells, and 5.22 µM (resistant) vs. 5.93 µM (parental) in 22Rv1 cells.

Stable FKBP7 knockdown has previously been shown to chemosensitize docetaxel-resistant IGR-CaP1-Rvivo xenografts *in vivo (12)*. Immunoblots showed that FKBP7 knockdown (sh5 and sh7) markedly reduced NFE2L1 levels without affecting NFE2L2 levels in these resistant cells (Fig. 6B), indicating that FKBP7 controls NFE2L1 levels but not NFE2L2 levels. Transient FKBP7 knockdown also slightly decreased NFE2L1 protein levels in IGR-CaP1 resistant cells (Fig. 6C). In contrast, complete depletion of NFE2L1 did not consistently affect FKBP7 protein levels, suggesting that NFE2L1 does not regulate FKBP7 expression (Fig. 6C). We also overexpressed wild-type WT-FKBP7 and the Phe137Tyr point mutant (F137Y) with an inactivated PPIase domain. The Phe137 residue in FKBP7 is analogous to Phe99 in the rapamycin pocket of FKBP12 (Fig. 4G and Fig. S10) and has been shown to be required for PPIase activity *(49, 50)*. In 22Rv1 cells, overexpression of WT-FKBP7 increased NFE2L1 levels in a dose dependent manner, whereas overexpression of the F137Y mutant did not (Fig. 6D), suggesting a role for FKBP7 PPIase activity in the regulation of NFE2L1 levels.

Both FKBP7 and NFE2L1 silencing reduced the proliferation of taxane-resistant cells (Fig. 6E), consistent with NFE2L1 acting downstream of FKBP7 in docetaxel resistance. Disrupting NFE2L1 activity by inhibiting NGLY1-mediated de-N-glycosylation of its cytoplasmic form with the WRR139 inhibitor *(51)* was highly effective in suppressing the growth of the two docetaxel-resistant cell lines, highlighting the importance of NFE2L1 for the survival of resistant cells (Fig. 6F). Lastly, we compared the efficiency of WRR139-mediated NFE2L1 inhibition between docetaxel-resistant and parental cell lines, and observed a slight improvement in response to WRR139 in resistant cells (IC_50_ 1.72 vs. 1.98µM in IGR-CaP1 cells; IC_50_ 5.22 vs. 5.93µM in 22Rv1 cells), indicating that resistant cells are more dependent on NFE2L1 than sensitive cells (Fig. 6G). Overall, these results demonstrate that FKBP7 positively regulates NFE2L1 levels, suggesting that high FKBP7 levels in resistant cells may contribute to NFE2L1-mediated cell survival.

## DISCUSSION

A detailed understanding of the mechanisms behind chemoresistance in PCa is required to develop new biologically driven treatments that can target or bypass them and improve patient survival. In this context, we have previously reported that FKBP7 interacts with the translation initiation factor eIF4G, indicating a potential role for FKBP7 in regulating translation to promote cell survival. Based on this work, we proposed FKBP7 as a potential therapeutic target in PCa cells resistant to taxane therapy *(12)*. However, little data on FKBP7 is available to date, and a deeper understanding of how it functions at the molecular and cellular levels is required for proper future targeting of resistant cancer cells. FKBP7 is expressed in the lumen of the endoplasmic reticulum and exhibits *cis*-*trans* prolyl isomerase activity. Here, we report for the first time that FKBP7 is refluxed into the cytosol where it undergoes a gain of function. This reflux, which occurs at low levels under native conditions, is increased by chemotherapy and in taxane-resistant cells, independently of the PCa cell line studied (Fig. 7). We observed that chemotherapy induces both oxidative stress and FKBP7 overexpression, which is associated with increased cytosolic FKBP7 levels, suggesting that FKBP7 cytosolic reflux could be an adaptive response to counteract this stress. In the cytosol, FKBP7 binds eIF4G1. The increase in FKBP7 cytosolic levels, particularly upon chemotherapy exposure or in chemoresistant cells, must increase the amount of FKBP7/eIF4G1 complexes. In addition to eIF4G1, we identified the 40S small ribosomal subunit as a new cytosolic partner of FKBP7, further supporting its involvement in translation. We demonstrated that inhibiting FKBP7 in docetaxel-resistant cells affects this process by buffering the translation of a large set of mRNAs. Many of these mRNAs are targets of the transcription factor NFE2L1, which is involved in adaptive responses to oxidative stress *(42)*. Our findings therefore provide evidence that FKBP7 participates in a translational regulatory mechanism contributing to adaptive resistance to taxanes in PCa, involving eIF4G1 and NFE2L1 and thought to enable resistant cells to withstand oxidative stress.

**Figure 7:**
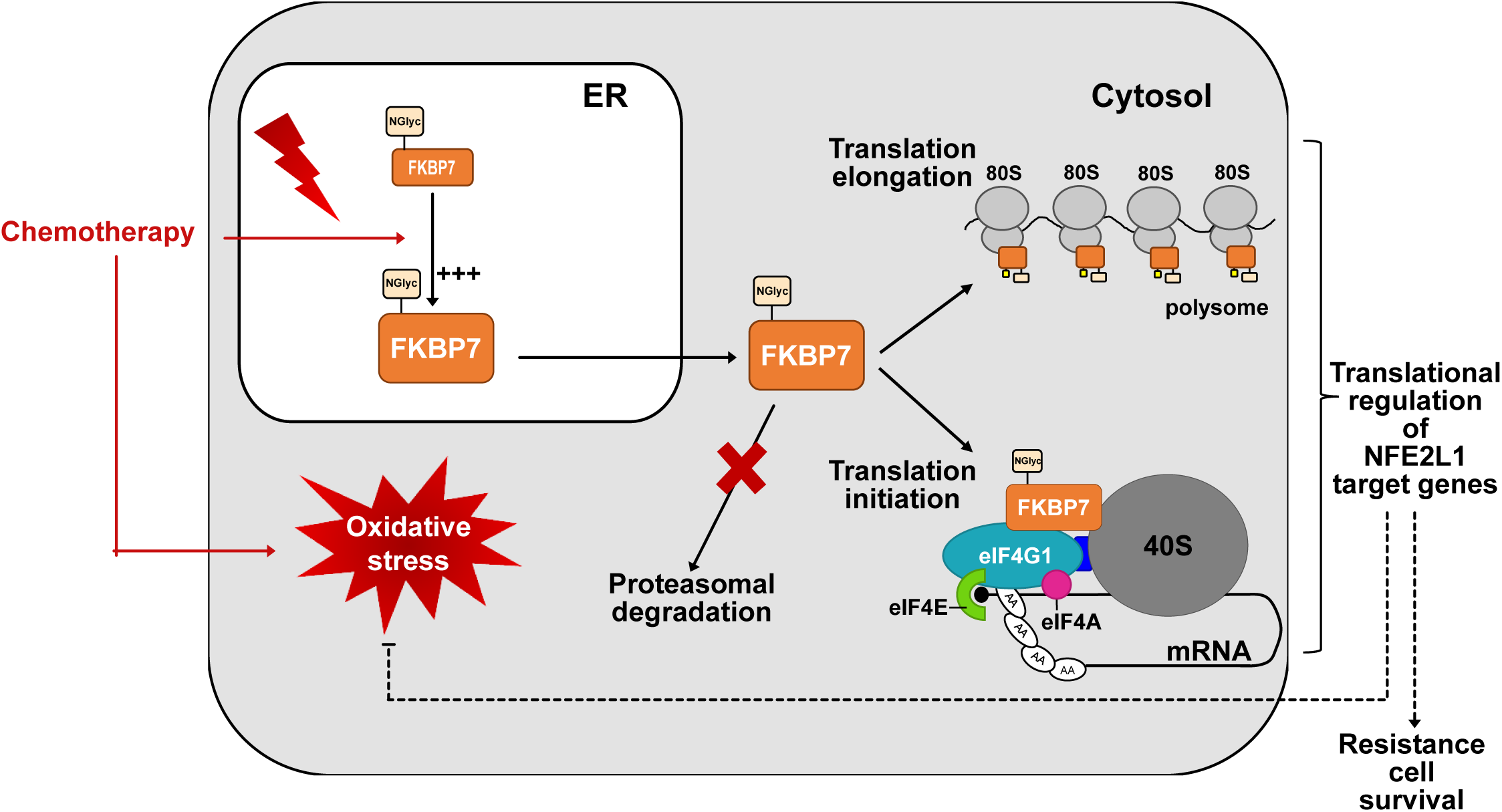
Proposed model for FKBP7 mechanism in docetaxel-resistant cells. As a luminal ER-resident protein, newly synthesized FKBP7 is N-glycosylated at position N45 in the ER. ER-levels of FKBP7 increased upon chemotherapy treatment, and the resulting oxidative stress enhances FKBP7 retrotranslocation into the cytosol, where the N-glycosylated form escapes ERAD-mediated proteasomal degradation. Once in the cytosol, FKBP7 acquires a moonlighting function in translation initiation, through binding to eIF4G1 and the 40S ribosomal subunit, and in translation elongation, through binding to polysomes. Based on the effects of FKBP7 silencing, cytosolic FKBP7 may contribute to the translation of NFE2L1 and other mRNAs involved in the oxidative stress response, thereby promoting the survival of docetaxel-resistant cells.

Cytosolic FKBP7 is not directly synthetized in the cytosol but rather refluxed from the ER, as it predominantly exists in an N-glycosylated form acquired in the ER. This glycosylated cytosolic form is not targeted by ER-associated degradation (ERAD), as its levels remain stable in the presence of a proteasome inhibitor. A short-lived, non-glycosylated form of FKBP7 was also detected at lower levels in the cytosol, whose role remains unknown but is presumably related to the ERAD. Non-degradative retrotranslocation of ER proteins to the cytosol has previously been described for calreticulin *(52)*, and for the protein disulfide isomerase AGR2/PDIA17 in glioblastoma cells *(6)*, where it operates as an active protein-dependent process. In the case of AGR2, this constitutively active process, termed ERCYS and demonstrated using orthogonal methods, acts as a pro-survival mechanism in response to ER stress, enhancing the oncogenic properties of cancer cells *(6)*. A recent spatial proteomics study in yeast further identified Fpr2, the yeast ER-resident FKBP homolog, among proteins relocating to the cytosol under ER stress *(53)*. While a passive leakage of FKBP7 from the ER to the cytosol under stress cannot be formally excluded, our results showed that cytosolic FKBP7 is associated with cytosolic ribosomal complexes, and not with ER-associated ribosomes, consistent with the possibility that this reflux corresponds to an ERCYS-like mechanism enabling cells to adapt to chemotherapy-induced stress.

Oxidative stress has emerged as a mechanism of drug resistance in cancer cells *(4)*. Chemotherapeutic drugs disrupt redox homeostasis by inducing large amounts of reactive oxygen species (ROS). Cancer cells can withstand excessive ROS levels by producing protective molecules, and metabolites generated during the resulting oxidative stress can favor cell survival and confer resistance to chemotherapy *(54)*. Here, we showed that cytosolic FKBP7 levels increased upon treatment with chemotherapeutic agents known to induce oxidative stress, such as the anthracycline doxorubicin and the topoisomerase inhibitor etoposide, both strong ROS inducers. Like other cytotoxic agents, docetaxel disrupted redox homeostasis by increasing both ROS levels and cytosolic FKBP7 levels. In contrast, docetaxel-resistant cells overexpressing FKBP7 appeared to have restored their redox homeostasis and no longer produce ROS under treatment. These results suggest that cytosolic FKBP7 may contribute to an early escape mechanism from chemotherapy-induced oxidative stress. Taxane-resistant cells, characterized by high FKBP7 levels, may exploit this process to cope with oxidative stress, maintain redox homeostasis, and escape cell death.

Translational reprogramming at the initiation and elongation steps of translation enables some tumor cells to evade the effects of drugs. This translational plasticity involves components of the eIF4F translation initiation complex, which have been shown to mediate resistance to several chemotherapeutic agents across cancers types *(8)*. Among the three eIF4G isoforms, eIF4G1 is amplified or overexpressed in multiple cancers and plays a vital role in cancer cell survival *(33)*, including in castration-resistant prostate cancer (CRPC) *(34)*. Having previously reported that FKBP7 interacts with eIF4G in docetaxel-resistant cells *(12)*, we identified two major interacting domains on eIF4G1, the C-terminal HEAT3 domain and the sequences flanking the HEAT1 domain, but not the HEAT1 domain in isolation. The C-terminal HEAT3 domain contains proline 1497 in a *cis* conformation, which might be stabilized by FKBP7 isomerization activity. This intermolecular interaction could therefore potentially contribute to the correct folding and cellular stabilization of eIF4G1, as previously observed *(12)*. Further structural data are needed to understand how the HEAT3 motif folds within the context of the full-length protein and to determine the accessibility of this proline residue and isomerization by FKBP7. Our data also reveal a previously unknown interaction between FKBP7 and RPS6, a component of the 40S small ribosomal subunit. Given that FKBP7 also interacts with eIF4G1, this raise the possibility that FKBP7 is present within the 48S initiation complex, although it may act at other steps of translation initiation as well. Consistently, we also found FKBP7 associated with polysomes, suggesting a role in translation elongation. FKBP10, another ER-resident FKBP, has similarly been detected in the cytosol *(6)*, and implicated in translation elongation *(16)*, suggesting a common regulatory mechanism involving several ER-resident FKBPs in cytosolic translation. Further support for this concept comes from other species, such as *Pf*FKBP35, the sole FKBP homolog in *Plasmodium falciparum*, which has also been implicated in translational regulation *(55)*.

In the present study, FKBP7 silencing in resistant cells reduced cell proliferation and triggered translational buffering of a large set of mRNAs. A similar translational offsetting process has been reported in PCa cells with transcriptional defects induced by estrogen receptor α depletion *(56)* and in yeast cells under acute oxidative stress *(57)*, where ribosome occupancy of the buffered mRNAs is adjusted to counteract changes in mRNA abundance and limit variation at the protein level. Although the mechanisms underlying translational buffering remain incompletely understood, this process is thought to play a key role in shaping proteome composition as an adaptive mode of gene expression regulation. We found that FKBP7 silencing-induced translational buffering was, at least in part, associated with modulation of gene transcriptionally regulated by NFE2L1, recently shown to be essential for maintaining redox homeostasis in adaptive responses to pathophysiological stress *(42)*, and implicated in metabolic reprogramming and ferroptosis resistance independently of NRF2/NFE2L2 *(58)*. We showed that FKBP7 silencing reduces NFE2L1 protein levels, whereas FKBP7 overexpression moderately increases them in an isomerase-dependent process. Together, these results suggest that FKBP7 could control NFE2L1 levels, thereby maintaining an essential protein balance under chemotherapy-induced stress. Regarding the underlying mechanism, FKBP7 could affect NFE2L1 through translational regulation, given its interaction with eIF4G1 and the large overlap between the FKBP7-dependent buffering gene set and NFE2L1 target genes, although our anota2seq analysis did not detect a direct effect on NFE2L1 translation at the 48-hour timepoint analyzed, possibly reflecting an early, transient regulatory event no longer detectable at this stage. We cannot rule out that FKBP7 may also affect NFE2L1 protein stability or activity independently of translation. In this study, taxane-resistant cells display FKBP7-driven increases in NFE2L1 levels, which may contribute to the metabolic reprogramming critical for resistant cell survival. Our GSEA analysis of the translationally buffered gene set revealed enrichment in fatty acid metabolism pathways (Fig. S8B), although the specific downstream targets of NFE2L1 within this gene set remain to be formally identified. Finally, targeting NFE2L1/NGLY1 with the WRR139 inhibitor efficiently overcame taxane resistance. As late-stage PCa is particularly subject to reprogramming of lipid metabolism and ferroptosis *(59)*, our results point to new potential therapeutic targets for taxane resistance in PCa.

As a first step towards drug design, we also presented the first structural and binding data on FKBP7. Using NMR, we showed that the FKBP7 PPIase domain is properly folded and binds to rapamycin and everolimus. However, unlike most human FKBP proteins *(60)*, FKBP7 did not interact with FK506, highlighting its distinct binding properties. The divergent residues at the ligand-binding site between FKBP7 and FKBP12 (Fig. 4G), including a Try-to-Leu substitution at position 59 (FKBP12 numbering), a Asp-to-Tyr substitution at position 37, or an Phe-to-Lys substitution at position 46, are potential determinants of these distinct FKBP7 binding properties and offer promising starting points for the future design of ligand selective for FKBP7 over other FKBPs.

In conclusion, we report an unexpected gain of function of the ER-resident protein FKBP7, which is overexpressed and refluxed into the cytosol upon chemotherapy, concomitant with increased oxidative stress. In the cytosol, FKBP7 interacts with eIF4G1 and ribosomal subunits, and its silencing buffers the translation of a large set of NFE2L1-dependent mRNAs. FKBP7 sustains NFE2L1 levels, a transcription factor known to promote redox homeostasis and cell survival. For future clinical applications, we present the first data on the ligand-binding specificities of the FKBP7 catalytic domain. Our study identifies the FKBP7/NFE2L1 axis as a promising target for the development of drugs to prevent chemoresistance in prostate cancer. As the regulation of FKBP7 levels is not restricted to taxanes but extends to a wide range of chemotherapeutic agents, targeting the FKBP7-NFE2L1 pathway could benefit the treatment of other solid tumors currently managed with chemotherapy.

## MATERIALS AND METHODS

### Cell lines

The human prostate cancer (PCa) cell lines and their drug-resistant derivatives IGR-CaP1, PC3, and 22Rv1 *(12)*, LNCaP *(61)* were maintained in RPMI 1640 medium with 10% fetal bovine serum (Sigma-Aldrich). Taxane-resistant populations were generated by long-term dose-escalating exposure to docetaxel, as previously described *(62)* and were maintained by monthly treatment with the maximum tolerated docetaxel dose. LNCaP derivatives 49F and 42D were cultured with 10 µM enzalutamide. The cells were regularly tested for the potential presence of mycoplasma (Eurofins Genomics).

### Reagents and Antibodies

MG132, rapamycin, everolimus and FK506 (Selleckchem), tunicamycin, thapsigargin and brefeldin A (Sigma-Aldrich), and cabazitaxel, enzalutamide, HA15 and WRR139 inhibitors (MedChemExpress) were resuspended in DMSO. Injectable chemotherapy solutions were purchased from Accord (docetaxel, etoposide, 5FU, and carboplatin) and from Mylan (methotrexate) and were resuspended in culture medium. Antibodies used in the study were described in table S1.

### Transfections and small interfering RNAs (siRNAs)

Depending on the experiment, cells were seeded 24 h before transfection or transfected at the time of seeding. For FKBP7 silencing, cells at ∼70% confluence were transiently transfected for 48 h with 20 nM stealth siRNA or Silencer siRNA targeting FKBP7 (table S1, ThermoFisher) using JetPrime reagent (PolyPlus) according to the supplier’s protocol. Silencer siRNAs (ThermoFisher) were similarly used to silence NFE2L1 and NFE2L2 (table S1), and the corresponding non-targeting control siRNA (siNT, ThermoFisher) was used under the same conditions. For coimmunoprecipitation experiments overexpressing FKBP7 and/or eIF4G1, cells were seeded 24 h before transfection to reach ∼70% confluence, and expression vectors were transfected for 48 h using JetOptimus reagent (PolyPlus), according to the supplier’s protocol.

### Immunoprecipitation (IP)

After transient transfection, cells were rinsed twice in cold PBS and lysed for 30 min at 4°C in lysis buffer (120 mM NaCl, 20 mM HEPES, 1 mM EDTA, 5 % glycerol, 0.5 % igepal, and protease and phosphatase inhibitors (ThermoFisher)). Supernatants were collected by centrifugation at 16,100 x g for 10 min at 4°C. Ten percent of the extract was kept as input, and the remainder was divided equally for a 2h-incubation at 4 °C with 1-1.5 µg of anti-FKBP7, anti-IgG, or anti-GFP antibody and magnetic beads (ThermoFisher). After three washes in lysis buffer, immunoprecipitated complexes were eluted, and complexes and inputs were denatured at 70°C for 10 min with LDS and reducing agent (ThermoFisher) before SDS-PAGE analysis.

### Subcellular fractionation and western blotting

Whole-cell lysates were prepared in RIPA buffer (Cell signaling Tech.) with protease and phosphatase inhibitors (ThermoFisher). Cytosolic and membrane fractions were isolated as previously described *(63)*. Briefly, after trypsinization and washing in PBS, cells were lysed for 10 min at 4°C with rotation in buffer containing 150 mM NaCl, 50 mM HEPES (pH 7.4), 25 µg/mL digitonin, and protease and phosphatase inhibitors, then centrifuged at 2000 x g for 5 min at 4°C; the supernatant constituted the cytosolic fraction. The pellet was washed with PBS and lysed for 30 min on ice in an equivalent volume of a buffer containing 150 mM NaCl, 50mM HEPES (pH 7.4), 1 % igepal, and protease and phosphatase inhibitors (ThermoFisher), then centrifuged at 7,400 x g at 4 °C for 10 min. The resulting supernatant constituted the membrane fraction. For immunoblotting, protein lysates were denatured in LDS buffer with reducing agent (ThermoFisher), separated by SDS-PAGE, and transferred onto a nitrocellulose membrane. After blocking with 5 % nonfat milk or BSA in TBS with 0.1 % Tween-20, membranes were incubated overnight with the primary antibodies (table S1). Proteins were detected with horseradish-peroxidase-conjugated secondary antibodies and enhanced chemiluminescence (Millipore) and imaged with the ChemiDoc XRS+ system (Bio-Rad Laboratories) using Image Lab software.

### Deglycosylation by enzymatic digestion

Cells were lysed by five 10-sec sonication pulses in 20 mM Tris-HCl (pH 7.5), 150 mM NaCl buffer supplemented with phosphatase inhibitors (ThermoFisher). Extracts were denatured at 100 °C for 10 min in the enzyme-specific denaturing buffer, then digested for 1h at 37°C with EndoH or PNGase F (both from New England Biolabs). Digestion products were analyzed by SDS-PAGE.

### NMR analysis

NMR experiments were performed on the purified FKBP7 PPIase domain (see Supplementary Materials for details) using a Bruker NEO 700 MHz spectrometer equipped with a TXO cryoprobe and an 800 MHz Avance III spectrometer equipped with a TCI cryoprobe. ^15^N SOFAST-HMQC spectra *(64)* were collected at 298 K on samples containing 50-200 µM ^15^N labeled FKBP7 in 50 mM HEPES (pH 7), 150 mM NaCl, 2 mM TCEP, 2 mM EDTA and 5% D_2_O. For titration experiments, DMSO-d6 stock solutions of rapamycin, everolimus and FK506 (5-10 mM) were added at the desired final concentration. As a control, DMSO-d6 was added to the protein solution at the desired amount. NMR spectra were processed with TopSpin and analyzed with CcpNmr (v2.5) *(65)*.

### Cell viability and cell proliferation assays

Cells were seeded in triplicate into 96-well plates at 3,500-10,000 cells per well. Cell viability was determined using a colorimetric WST-1-based assay (Roche) after 48-72 h of drug treatment and compared to untreated cells to calculate the surviving fraction. Dose-response curves and IC50 were estimated by weighted four-parameter logistic regression in GraphPad Prism (v10.4.1). Cell proliferation was determined by measuring cell confluency by live-cell time-lapse imaging Incucyte (Sartorius) after drug treatment or after reverse transfection with 20 nM siRNA.

### Reactive oxygen species (ROS) measurement

Overall cellular ROS levels were assessed with 2′,7′-dichlorofluorescein diacetate (H_2_DCFDA, ThermoFisher). Cells were seeded in triplicate at 3,500-10,000 cells/well, depending on the cell line, into black-walled, clear-bottom 96-well microplates (PerkinElmer) one day before the experiment. After 24 h, cells were treated with increasing doses of chemotherapy for 48 h. Cells were then washed once with serum-free medium and incubated with 10 µM H_2_DCFDA in serum-free medium for 30 min at 37°C. The H_2_DCFDA-containing supernatant was then removed, cells were washed twice, and culture medium was added. Green fluorescence and cell confluence were immediately measured with the Incucyte Live-Cell Analysis System (Sartorius). Results were expressed as total green fluorescence integrated intensity object average (GCU x µm^2^) relative to confluence (%) for each triplicate, with untreated cells as a reference condition.

### Glutathione measurements

Cells were seeded in triplicate at 3,500-10,000 cells/well, depending on the cell line in 96-well plates and grown overnight before treatment with chemotherapy for 48 hours. Cell confluence was determined in each well with the Incucyte Live-Cell Analysis System (Sartorius), and the glutathione levels were quantified with the GSH-Glo Glutathione Assay Kit (Promega, V6912), following the manufacturer’s instructions. Luminescence was measured with a Spark microplate reader (Tecan). Glutathione levels were normalized to cell confluence before calculating the GSH/GSSG ratio.

### Polysome profiling

Sucrose density gradient centrifugation was used to separate the sub-polysomal and polysomal ribosome fractions, essentially as previously described *(66)*. Briefly, cells at ∼70% confluency were treated with 100 µg/ml cycloheximide (Sigma-Aldrich) for 5 min at 37°C, collected in cold PBS containing 100 µg/ml cycloheximide at the same concentration, centrifuged at 350 x g for 6 min at 4 °C, and resuspended in 500 µl of hypotonic buffer (5 mM Tris (pH 7.5), 2.5 mM MgCl_2_, 1.5 mM KCl, 100 µg/ml cycloheximide, 200 µM DTT) supplemented with RNase inhibitor (Invitrogen) and phosphatase and protease inhibitors (Roche). After homogenization, the lysate was adjusted to 0.5 % Triton X-100 and 0.5 % sodium deoxycholate, kept on ice for 30 min, and centrifuged at 16,100 x g for 10 min at 4 °C. RNA concentration in the resulting lysate was determined and adjusted to 260 µg/ml with hypotonic buffer. Ten percent of the lysate was kept as input, and the remainder was loaded onto a 5-50% sucrose gradient and centrifuged at 36,000 rpm (SW41 Ti rotor, Beckman) for 2 h at 4 °C. Polysome fractions were monitored and collected using either a gradient fractionation system (Isco, absorbance at 254 nm) or a Biocomp fractionator (absorbance at 260 nm). For high-throughput sequencing processing, input RNA and polysome-bound RNA (pool of the last five polysome fractions) were extracted using Trizol LS (ThermoFisher) and quantified on NanoDrop 2000 (Thermo Scientific). Each condition was performed in three independent experiments.

To specifically isolate cytosolic ribosomal complexes and exclude potential contamination by ER-associated membranes, a variant of this protocol was used to extract ribosomes from the cytoplasmic fraction, based on selective digitonin permeabilization of the plasma membrane *(67)*. Cells were treated with cycloheximide and harvested as described above, then resuspended in cytosolic extraction buffer (25 mM HEPES-KOH pH 7.4, 100 mM KCl, 5 mM MgCl2, 1 mM DTT, 100 µg/ml cycloheximide, RNase inhibitor, EDTA-free protease inhibitor cocktail, 0.03% digitonin) for 5 min on ice with gentle mixing; this low digitonin concentration selectively permeabilizes the plasma membrane while minimizing solubilization of the ER *(67)*. Lysates were cleared by sequential centrifugation (2,000 × g, 5 min, then 12,000 × g, 5 min, 4°C), and an aliquot of the clarified cytosolic fraction was kept for western blot analysis of cytosolic, ribosomal, and ER markers. Ribosome-containing complexes were pelleted by ultracentrifugation of the clarified fraction (100,000 × g, 2 h, 4°C) in the same buffer, then resuspended in digitonin-free buffer and split into two aliquots: one untreated and one supplemented with EDTA (30 mM, 30 min on ice) to chelate Mg2+ and dissociate 80S ribosomes and polysomes into 40S/60S subunits. Both aliquots were then loaded onto a sucrose gradient and processed as described above.

### Whole-transcriptome RNA-seq

The RNA integrity (RNA Integrity Score ≥ 7.0) was checked with the Fragment Analyzer (Agilent), and quantity was determined with Qubit (Invitrogen). Libraries were prepared with the SureSelect Automated Strand-Specific RNA Library Preparation Kit (Agilent) on the Bravo Platform (Agilent), according to the manufacturer’s instructions. Briefly, 50-200 ng of total RNA per sample was used for poly-A mRNA selection using Oligo(dT) beads and thermal mRNA fragmentation, followed by cDNA synthesis and conversion into double-stranded DNA using kit reagents; the resulting dsDNA was used for library preparation. Libraries were barcoded, purified, pooled together at equal concentrations, and sequenced in paired-end mode (2 x 100 bp) on a NovaSeq 6000 (Illumina) at the Gustave Roussy genomics facility. RNA-seq read processing and translatome analysis with anota2seq are described in detail in the Supplementary Materials.

### Quantitative real-time RT-PCR

Total RNA was extracted from the cells using TRIzol reagent (Invitrogen), according to the manufacturer’s protocol, and quantified using a NanoDrop Spectrophotometer. One microgram of RNA was reverse-transcribed into cDNA using SuperScript II Reverse Transcriptase (Invitrogen). The resulting cDNA was amplified with Absolute Blue Master Mix (ThermoFischer) and gene-specific forward and reverse primers (0.4 µM) for 40 cycles on a ViiA 7 Real-Time PCR System (Applied Biosystems). Primers are listed in table S1. Relative gene expression was calculated using the 2^-ΔΔCt method, using TBP as the reference gene. Data represent the mean ± SD of three independent experiments.

### Statistical Analysis

Paired or unpaired Student’s t-test for two comparisons, one-way or two-way ANOVA for multiple comparisons followed by Dunnett’s post hoc test were applied in GraphPad Prism (v10.5) with P ≤ 0.05 considered significant. *: P < 0.05; **: P < 0.01; ***: P < 0.001; ****: P < 0.0001; ns: not significant.

## DATA AVAIBILITY

The RNA-seq datasets generated during this study have been deposited in the Gene Expression Omnibus (GEO) database under the accession number GSE322750. Scripts and pipeline are available on Zenodo repository accessible at https://doi.org/10.5281/zenodo.XXXXXX. The raw NMR spectral data have been deposited on Zenodo at https://doi.org/10.5281/zenodo.XXXXXX. The plasmids and taxane-resistant cell lines generated in this study are available upon request or under a material transfer agreement. All other data needed to evaluate the conclusions in the paper are present in the paper or the Supplementary Materials.

## ACKNOWLEDGEMENTS AND FUNDING SOURCES

We thank S. Pyronnet (Cancer Research Center, Toulouse) for the kind gift of the eIF4G1 expression vectors, O. Larsson and I. Aksoylu (Karolinska Institute, Stockholm) for their advice in polysome profiling data analysis, and M. Tannoury and S. Lacoste for experimental help. This work was performed thanks to Gustave Roussy core facilities (Genomics and Bioinformatics). It has been supported by INSERM, the Fondation ARC pour la recherche sur le cancer (PJA20181208245), the Taxe d’Apprentissage LUDR2020, the PARRAINAGE CHERCHEUR charity program from Gustave Roussy, the French Infrastructure for Integrated Structural Biology (FRISBI) ANR-10-INBS-0005 and through the TRANSLACORE COST Action (CA21154). Financial support from the IR INFRANALYTICS FR2054 for conducting the research is gratefully acknowledged. LD received support from the Ministère de l’Enseignement Supérieur et de la Recherche and the Fondation ARC, AA-L received support from France Excellence Eiffel. FO and BC were supported by the PARRAINAGE CHERCHEUR charity program.

## Preprint

BioRxiv preprint V1 doi: https://doi.org/10.64898/2026.01.18.700172; this version posted January 19, 2026.

## REFERENCES

1. C. L. Vale, S. Burdett, L. H. M. Rydzewska, L. Albiges, N. W. Clarke, D. Fisher, K. Fizazi, G. Gravis, N. D. James, M. D. Mason, M. K. B. Parmar, C. J. Sweeney, M. R. Sydes, B. Tombal, J. F. Tierney, Addition of docetaxel or bisphosphonates to standard of care in men with localised or metastatic, hormone-sensitive prostate cancer: a systematic review and meta-analyses of aggregate data. The Lancet. Oncology 17, 243– 256 (2016).

2. M. Carceles-Cordon, V. Rodriguez-Bravo, D. P. Petrylak, J. Domingo-Domenech, 20 years of taxane therapy in prostate cancer - the past, present and future. Nature reviews. Urology (2025), doi:10.1038/S41585-025-01104-9.

3. W. Zhang, Y. Shi, L. Oyang, S. Cui, S. Li, J. Li, L. Liu, Y. Li, M. Peng, S. Tan, L. Xia, J. Lin, X. Xu, N. Wu, Q. Peng, Y. Tang, X. Luo, Q. Liao, X. Jiang, Y. Zhou, Endoplasmic reticulum stress-a key guardian in cancer. Cell Death Discov 10, 343 (2024).

4. X. Liang, J. Weng, Z. You, Y. Wang, J. Wen, Z. Xia, S. Huang, P. Luo, Q. Cheng, Oxidative stress in cancer: from tumor and microenvironment remodeling to therapeutic frontiers. Molecular cancer 24 (2025), doi:10.1186/S12943-025-02375-X.

5. E. Chevet, C. Hetz, A. Samali, Endoplasmic reticulum stress-activated cell reprogramming in oncogenesis. Cancer discovery 5, 586–97 (2015).

6. D. Sicari, F. G. Centonze, R. Pineau, P. Le Reste, L. Negroni, S. Chat, M. A. Mohtar, D. Thomas, R. Gillet, T. Hupp, E. Chevet, A. Igbaria, Reflux of Endoplasmic Reticulum proteins to the cytosol inactivates tumor suppressors. EMBO reports 22 (2021), doi:10.15252/EMBR.202051412.

7. D. D. Zhang, Thirty years of NRF2: advances and therapeutic challenges. Nature Reviews Drug Discovery 24, 421–444 (2025).

8. L. Fabbri, A. Chakraborty, C. Robert, S. Vagner, The plasticity of mRNA translation during cancer progression and therapy resistance. Nature reviews. Cancer 21, 558–577 (2021).

9. M. Bhat, N. Robichaud, L. Hulea, N. Sonenberg, J. Pelletier, I. Topisirovic, Targeting the translation machinery in cancer. Nature reviews. Drug discovery 14, 261–78 (2015).

10. S. Joshi, T. Wang, T. L. S. Araujo, S. Sharma, J. L. Brodsky, G. Chiosis, Adapting to stress — chaperome networks in cancer. Nat Rev Cancer 18, 562–575 (2018).

11. C. Zhang, J. Li, Q. Tang, L. Li, D. Cao, Targeting proteostasis for cancer therapy: current advances, challenges, and future perspectives. Mol Cancer 24, 265 (2025).

12. M. F. Garrido, N. J. P. Martin, M. Bertrand, C. Gaudin, F. eric Commo, N. E. Kalaany, N. A. Nakouzi, L. Fazli, E. D. Nery, J. Camonis, F. Perez, S. Lerondel, A. Le Pape, D. Compagno, M. Gleave, Y. Loriot, L. Desaubry, S. Vagner, K. Fizazi, A. Chauchereau, Regulation of eIF4F Translation Initiation Complex by the Peptidyl Prolyl Isomerase FKBP7 in Taxane-resistant Prostate Cancer. Clinical Cancer Research 25, 710–723 (2019).

13. S. Romano, A. D’Angelillo, M. F. Romano, Pleiotropic roles in cancer biology for multifaceted proteins FKBPs. Biochimica et biophysica acta 1850, 2061–8 (2015).

14. J. M. Bonner, G. L. Boulianne, Diverse structures, functions and uses of FK506 binding proteins. Cellular signalling 38, 97–105 (2017).

15. F. neng Jiang, L. jun Dai, S. bang Yang, Y. ding Wu, Y. xiang Liang, X. li Yin, C. yun Zou, W. de Zhong, Increasing of FKBP9 can predict poor prognosis in patients with prostate cancer. Pathology Research and Practice 216 (2020), doi:10.1016/j.prp.2019.152732.

16. G. Ramadori, R. M. Ioris, Z. Villanyi, R. Firnkes, O. O. Panasenko, G. Allen, G. Konstantinidou, E. Aras, X. Brenachot, T. Biscotti, A. Charollais, M. Luchetti, F. Bezrukov, A. Santinelli, M. Samad, P. Baldi, M. A. Collart, R. Coppari, FKBP10 Regulates Protein Translation to Sustain Lung Cancer Growth. Cell Rep 30, 3851–3863.e6 (2020).

17. S. Oh, J. Yeom, H. J. Cho, J. H. Kim, S. J. Yoon, H. Kim, J. K. Sa, S. Ju, H. Lee, M. J. Oh, W. Lee, Y. Kwon, H. Li, S. Choi, J. H. Han, J. H. Chang, E. Choi, J. Kim, N. G. Her, S. H. Kim, S. G. Kang, E. Paek, D. H. Nam, C. Lee, H. S. Kim, Integrated pharmaco-proteogenomics defines two subgroups in isocitrate dehydrogenase wild-type glioblastoma with prognostic and therapeutic opportunities. Nature communications 11 (2020), doi:10.1038/S41467-020-17139-Y.

18. M. Hagedorn, G. Siegfried, K. B. Hooks, A. M. Khatib, Integration of zebrafish fin regeneration genes with expression data of human tumors in silico uncovers potential novel melanoma markers. Oncotarget 7, 71567–71579 (2016).

19. H. Kim, Y. Feng, R. Murad, J. Pozniak, C. Pelz, Y. Chen, B. Dalal, R. Sears, E. Sergienko, M. Jackson, E. Ruppin, M. Herlyn, C. Harris, J. C. Marine, V. Klepsch, G. Baier, Z. A. Ronai, Melanoma-intrinsic NR2F6 activity regulates antitumor immunity. Science advances 9 (2023), doi:10.1126/SCIADV.ADF6621.

20. M. Feng, C. Gu, S. Ma, Y. Wang, H. Liu, R. Han, J. Gao, Y. Long, H. Mi, Mouse FKBP23 mediates conformer-specific functions of BiP by catalyzing Pro117 cis/trans isomerization. Biochemical and biophysical research communications 408, 537–40 (2011).

21. Y. Ishikawa, K. Mizuno, H. P. Bächinger, Ziploc-ing the structure 2.0: Endoplasmic reticulum-resident peptidyl prolyl isomerases show different activities toward hydroxyproline. The Journal of biological chemistry 292, 9273–9282 (2017).

22. X. Zhang, Y. Wang, H. Li, W. Zhang, D. Wu, H. Mi, The mouse FKBP23 binds to BiP in ER and the binding of C-terminal domain is interrelated with Ca2+ concentration. FEBS Lett 559, 57–60 (2004).

23. Y. Wang, R. Han, D. Wu, J. Li, C. Chen, H. Ma, H. Mi, The binding of FKBP23 to BiP modulates BiP’s ATPase activity with its PPIase activity. Biochemical and biophysical research communications 354, 315–20 (2007).

24. T. Nakamura, D. Yabe, N. Kanazawa, K. Tashiro, S. Sasayama, T. Honjo, Molecular cloning, characterization, and chromosomal localization of FKBP23, a novel FK506-binding protein with Ca2+-binding ability. Genomics 54, 89–98 (1998).

25. J. J. Caramelo, A. J. Parodi, A sweet code for glycoprotein folding. FEBS Letters 589, 3379–3387 (2015).

26. L. Qi, B. Tsai, P. Arvan, New Insights into the Physiological Role of Endoplasmic Reticulum-Associated Degradation. Trends in cell biology 27, 430–440 (2017).

27. M. V. Blagosklonny, G. S. Wu, S. Omura, W. S. El-Deiry, Proteasome-dependent regulation of p21WAF1/CIP1 expression. Biochemical and biophysical research communications 227, 564–569 (1996).

28. T. Avril, E. Vauléon, E. Chevet, Endoplasmic reticulum stress signaling and chemotherapy resistance in solid cancers. Oncogenesis 6 (2017), doi:10.1038/ONCSIS.2017.72.

29. M. Cerezo, A. Lehraiki, A. Millet, F. Rouaud, M. Plaisant, E. Jaune, T. Botton, C. Ronco, P. Abbe, H. Amdouni, T. Passeron, V. Hofman, B. Mograbi, A.-S. Dabert-Gay, D. Debayle, D. Alcor, N. Rabhi, J.-S. Annicotte, L. Héliot, M. Gonzalez-Pisfil, C. Robert, S. Moréra, A. Virougoux, P. Gual, M. M. U. Ali, C. Bertolotto, P. Hofman, R. Ballotti, R. Benhida, S. Rocchi, Compounds Triggering ER Stress Exert Anti-Melanoma Effects and Overcome BRAF Inhibitor Resistance. Cancer cell 29, 805–19 (2016).

30. C. Ruggiero, M. Doghman-Bouguerra, C. Ronco, R. Benhida, S. Rocchi, E. Lalli, The GRP78/BiP inhibitor HA15 synergizes with mitotane action against adrenocortical carcinoma cells through convergent activation of ER stress pathways. Molecular and Cellular Endocrinology 474, 57–64 (2018).

31. C. Gorrini, I. S. Harris, T. W. Mak, Modulation of oxidative stress as an anticancer strategy. Nature Reviews Drug Discovery 12, 931–947 (2013).

32. A. Paschos, R. Pandya, W. C. M. Duivenvoorden, J. H. Pinthus, Oxidative stress in prostate cancer: Changing research concepts towards a novel paradigm for prevention and therapeutics. Prostate Cancer and Prostatic Diseases 16, 217–225 (2013).

33. S. Wu, G. Wagner, Deep computational analysis details dysregulation of eukaryotic translation initiation complex eIF4F in human cancers. Cell systems 12, 907–923.e6 (2021).

34. P. K. Jaiswal, S. Koul, P. S. T. Shanmugam, H. K. Koul, Eukaryotic Translation Initiation Factor 4 Gamma 1 (eIF4G1) is upregulated during Prostate cancer progression and modulates cell growth and metastasis. Scientific reports 8 (2018), doi:10.1038/S41598-018-25798-7.

35. J. Pelletier, N. Sonenberg, The Organizing Principles of Eukaryotic Ribosome Recruitment. Annu Rev Biochem 88, 307–335 (2019).

36. D. Friedrich, A. Marintchev, H. Arthanari, The metaphorical swiss army knife: The multitude and diverse roles of HEAT domains in eukaryotic translation initiation. Nucleic acids research 50, 5424–5442 (2022).

37. L. Bellsolell, P. F. Cho-Park, F. Poulin, N. Sonenberg, S. K. Burley, Two structurally atypical HEAT domains in the C-terminal portion of human eIF4G support binding to eIF4A and Mnk1. Structure (London, England : 1993) 14, 913–923 (2006).

38. L. A. Kelley, S. Mezulis, C. M. Yates, M. N. Wass, M. J. E. Sternberg, The Phyre2 web portal for protein modeling, prediction and analysis. Nature protocols 10, 845–858 (2015).

39. J. Jumper, R. Evans, A. Pritzel, T. Green, M. Figurnov, O. Ronneberger, K. Tunyasuvunakool, R. Bates, A. Žídek, A. Potapenko, A. Bridgland, C. Meyer, S. A. A. Kohl, A. J. Ballard, A. Cowie, B. Romera-Paredes, S. Nikolov, R. Jain, J. Adler, T. Back, S. Petersen, D. Reiman, E. Clancy, M. Zielinski, M. Steinegger, M. Pacholska, T. Berghammer, S. Bodenstein, D. Silver, O. Vinyals, A. W. Senior, K. Kavukcuoglu, P. Kohli, D. Hassabis, Highly accurate protein structure prediction with AlphaFold. Nature 596, 583–589 (2021).

40. S. P. Boudko, Y. Ishikawa, J. Nix, M. S. Chapman, H. P. Bächinger, Structure of human peptidyl-prolyl cis-trans isomerase FKBP22 containing two EF-hand motifs. Protein Science 23, 67–75 (2014).

41. C. Oertlin, J. Lorent, C. Murie, L. Furic, I. Topisirovic, O. Larsson, Generally applicable transcriptome-wide analysis of translation using anota2seq. Nucleic Acids Research 47, e70–e70 (2019).

42. X. Liu, C. Xu, W. Xiao, N. Yan, Unravelling the role of NFE2L1 in stress responses and related diseases. Redox Biology 65 (2023), doi:10.1016/j.redox.2023.102819.

43. M. B. Murray, S. J. Dixon, Ferroptosis regulation by Cap’n’collar family transcription factors. Journal of Biological Chemistry 300 (2024), doi:10.1016/j.jbc.2024.107583.

44. P. Liu, M. J. Kerins, W. Tian, D. Neupane, D. D. Zhang, A. Ooi, Differential and overlapping targets of the transcriptional regulators NRF1, NRF2, and NRF3 in human cells. Journal of Biological Chemistry 294, 18131–18149 (2019).

45. L. Ibrahim, J. Mesgarzadeh, I. Xu, E. T. Powers, R. Luke Wiseman, M. J. Bollong, Defining the functional targets of capŉ‘collar transcription factors NRF1, NRF2, and NRF3. Antioxidants 9, 1–15 (2020).

46. N. J. Lehrbach, P. C. Breen, G. Ruvkun, Protein Sequence Editing of SKN-1A/Nrf1 by Peptide:N-Glycanase Controls Proteasome Gene Expression. Cell 177, 737–750.e15 (2019).

47. K. P. Shay, A. J. Michels, W. Li, A. N. T. Kong, T. M. Hagen, Cap-independent Nrf2 translation is part of a lipoic acid-stimulated detoxification stress response. Biochimica et Biophysica Acta - Molecular Cell Research 1823, 1102–1109 (2012).

48. J. Steffen, M. Seeger, A. Koch, E. Krüger, Proteasomal degradation is transcriptionally controlled by TCF11 via an ERAD-dependent feedback loop. Molecular Cell 40, 147–158 (2010).

49. D. L. Riggs, M. B. Cox, H. L. Tardif, M. Hessling, J. Buchner, D. F. Smith, Noncatalytic Role of the FKBP52 Peptidyl-Prolyl Isomerase Domain in the Regulation of Steroid Hormone Signaling. Molecular and Cellular Biology 27, 8658–8669 (2007).

50. G. Gudavicius, H. Soufari, S. K. Upadhyay, S. K. Upadhyay, C. D. Mackereth, C. J. Nelson, Resolving the functions of peptidylprolyl isomerases: insights from the mutagenesis of the nuclear FKBP25 enzyme. Biochemical Society transactions 41, 761–8 (2013).

51. F. M. Tomlin, U. I. M. Gerling-Driessen, Y. C. Liu, R. A. Flynn, J. R. Vangala, C. S. Lentz, S. Clauder-Muenster, P. Jakob, W. F. Mueller, D. Ordoñez-Rueda, M. Paulsen, N. Matsui, D. Foley, A. Rafalko, T. Suzuki, M. Bogyo, L. M. Steinmetz, S. K. Radhakrishnan, C. R. Bertozzi, Inhibition of NGLY1 Inactivates the Transcription Factor Nrf1 and Potentiates Proteasome Inhibitor Cytotoxicity. ACS Central Science 3, 1143– 1155 (2017).

52. G. Petris, L. Vecchi, M. Bestagno, O. R. Burrone, Efficient detection of proteins retro-translocated from the ER to the cytosol by in vivo biotinylation. PloS one 6 (2011), doi:10.1371/JOURNAL.PONE.0023712.

53. A. Platzek, K. Odehnalová, J. P. Schessner, G. H. H. Borner, S. Schuck, Dynamic Organellar Mapping in yeast reveals extensive protein localization changes during ER stress. Nat Commun 16, 10842 (2025).

54. J. D. Hayes, A. T. Dinkova-Kostova, K. D. Tew, Oxidative Stress in Cancer. Cancer cell 38, 167–197 (2020).

55. B. T. Thommen, J. M. Dziekan, F. Achcar, S. Tjia, A. Passecker, K. Buczak, C. Gumpp, A. Schmidt, M. Rottmann, C. Grüring, M. Marti, Z. Bozdech, N. M. B. Brancucci, Genetic validation of Pf FKBP35 as an antimalarial drug target. eLife 12 (2023), doi:10.7554/ELIFE.86975.

56. J. Lorent, E. P. Kusnadi, V. van Hoef, R. J. Rebello, M. Leibovitch, J. Ristau, S. Chen, M. G. Lawrence, K. J. Szkop, B. Samreen, P. Balanathan, F. Rapino, P. Close, P. Bukczynska, K. Scharmann, I. Takizawa, G. P. Risbridger, L. A. Selth, S. A. Leidel, Q. Lin, I. Topisirovic, O. Larsson, L. Furic, Translational offsetting as a mode of estrogen receptor α-dependent regulation of gene expression. The EMBO journal 38 (2019), doi:10.15252/EMBJ.2018101323.

57. W. R. Blevins, T. Tavella, S. G. Moro, B. Blasco-Moreno, A. Closa-Mosquera, J. Díez, L. B. Carey, M. M. Albà, Extensive post-transcriptional buffering of gene expression in the response to severe oxidative stress in baker’s yeast. Scientific Reports 9 (2019), doi:10.1038/S41598-019-47424-W.

58. G. C. Forcina, L. Pope, M. Murray, W. Dong, M. Abu-Remaileh, C. R. Bertozzi, S. J. Dixon, Ferroptosis regulation by the NGLY1/NFE2L1 pathway. Proceedings of the National Academy of Sciences of the United States of America 119 (2022), doi:10.1073/PNAS.2118646119.

59. J. Liang, Y. Liao, P. Wang, K. Yang, Y. Wang, K. Wang, B. Zhong, D. Zhou, Q. Cao, J. Li, Y. Zhao, N. Jiang, Ferroptosis landscape in prostate cancer from molecular and metabolic perspective. Cell Death Discovery 9 (2023), doi:10.1038/S41420-023-01430-0.

60. J. Dornan, P. Taylor, M. D. Walkinshaw, Structures of immunophilins and their ligand complexes. Curr Top Med Chem 3, 1392–1409 (2003).

61. A. Zoubeidi, M. Gleave, Small heat shock proteins in cancer therapy and prognosis. International Journal of Biochemistry and Cell Biology 44, 1646–1656 (2012).

62. N. Al Nakouzi, S. Cotteret, F. Commo, C. Gaudin, S. Rajpar, P. Dessen, P. Vielh, K. Fizazi, A. Chauchereau, Targeting CDC25C, PLK1 and CHEK1 to overcome Docetaxel resistance induced by loss of LZTS1 in prostate cancer. Oncotarget 5, 667–78 (2014).

63. P. Holden, W. A. Horton, Crude subcellular fractionation of cultured mammalian cell lines. BMC Res Notes 2, 243 (2009).

64. P. Schanda, B. Brutscher, Very fast two-dimensional NMR spectroscopy for real-time investigation of dynamic events in proteins on the time scale of seconds. Journal of the American Chemical Society 127, 8014–8015 (2005).

65. W. F. Vranken, W. Boucher, T. J. Stevens, R. H. Fogh, A. Pajon, M. Llinas, E. L. Ulrich, J. L. Markley, J. Ionides, E. D. Laue, The CCPN data model for NMR spectroscopy: development of a software pipeline. Proteins 59, 687–696 (2005).

66. C. Cirotti, S. Rizza, P. Giglio, N. Poerio, M. F. Allega, G. Claps, C. Pecorari, J.-H. Lee, B. Benassi, D. Barilà, C. Robert, J. S. Stamler, F. Cecconi, M. Fraziano, T. T. Paull, G. Filomeni, Redox activation of ATM enhances GSNOR translation to sustain mitophagy and tolerance to oxidative stress. EMBO Rep 22, e50500 (2021).

67. D. W. Reid, C. V. Nicchitta, Primary role for endoplasmic reticulum-bound ribosomes in cellular translation identified by ribosome profiling. J Biol Chem 287, 5518–5527 (2012).

68. S. Pyronnet, H. Imataka, A. C. Gingras, R. Fukunaga, T. Hunter, N. Sonenberg, Human eukaryotic translation initiation factor 4G (eIF4G) recruits mnk1 to phosphorylate eIF4E. The EMBO journal 18, 270– 279 (1999).

69. S. Chen, Y. Zhou, Y. Chen, J. Gu, fastp: an ultra-fast all-in-one FASTQ preprocessor. *Bioinformatics (Oxford*, England) 34, i884–i890 (2018).

70. R. Patro, G. Duggal, M. I. Love, R. A. Irizarry, C. Kingsford, Salmon provides fast and bias-aware quantification of transcript expression. Nature methods 14, 417–419 (2017).

71. C. Soneson, M. I. Love, M. D. Robinson, Differential analyses for RNA-seq: Transcript-level estimates improve gene-level inferences. F1000Research 4 (2016), doi:10.12688/F1000RESEARCH.7563.2.

72. P. Danecek, J. K. Bonfield, J. Liddle, J. Marshall, V. Ohan, M. O. Pollard, A. Whitwham, T. Keane, S. A. McCarthy, R. M. Davies, Twelve years of SAMtools and BCFtools. GigaScience 10 (2021), doi:10.1093/GIGASCIENCE/GIAB008.

73. L. Wang, S. Wang, W. Li, RSeQC: quality control of RNA-seq experiments. *Bioinformatics (Oxford*, England) 28, 2184–2185 (2012).

74. S. W. Wingett, S. Andrews, FastQ Screen: A tool for multi-genome mapping and quality control. F1000Research 7, 1338–1338 (2018).

75. P. Ewels, M. Magnusson, S. Lundin, M. Käller, MultiQC: summarize analysis results for multiple tools and samples in a single report. *Bioinformatics (Oxford*, England) 32, 3047–3048 (2016).

76. J. Köster, S. Rahmann, Snakemake--a scalable bioinformatics workflow engine. *Bioinformatics (Oxford*, England) 28, 2520–2522 (2012).

