## Supplemental materials for "Cytoplasmic FKBP7 reflux sustains NFE2L1 levels in an adaptive response to chemotherapy in prostate cancer cells"

**SUPPLEMENTARY MATERIALS**

### Plasmid constructions

The 222 amino acids (aa) coding sequence for human FKBP7 (FKBP7-WT; NM_181342.3) was cloned into the pcDNA3.1 expression vector between the HindIII and XhoI sites. In all the FKBP7 expression vectors, nucleotides (nt) 513-537 were modified by site-directed mutagenesis (Q5 kit, New England Biolabs) to render it insensitive to siFKBP7-2 without altering the encoded protein. The FKBP7 N45Q mutant was generated by site-directed mutagenesis from FKBP7-WT, substituting Asn45 with Gln (codon 133-135; AAC to CAA), the F137Y mutant was generated by substituting Phe137 with Tyr (codon 409-411; TTT to TAT). The FLAG-tagged FKBP7 vector was designed from the predicted FKBP7 structure by inserting an internal FLAG tag after nucleotide 88 (i.e. after aa Glu29) via gene synthesis (Eurofins Genomics) and cloning into the HindIII/XhoI sites of the pcDNA3.1. To produce the FKBP7 catalytic domain recombinant protein, the sequence encoding amino acids 31-145, preceded by a TEV cleavage site, was obtained by gene synthesis (Eurofins Genomics) and cloned into the BamHI/HindIII sites of the His6-tag expression plasmid pET28b(+) (Merck Millipore).

Expression vectors encoding HA-tagged forms of the long eIF4G1 (NM_182917.4; HA-eIF4G1 long [aa 192-1600]) and the eIF4G1 fragments (HA-eIF4G1-N [aa 182-654]; HA-eIF4G1-M [aa 674-1079]; HA-eIF4G1-C [aa 1080-1600]) were provided by S. Pyronnet *(68)*. Expression vectors encoding HA-tagged forms of the eIF4G1 HEAT domains (4G1-HEAT1 [aa 752-990], 4G1-HEAT2 [aa 1235-1420], 4G1-HEAT3 [1438-1593], and 4G1-HEAT3-P1497A) were obtained by gene synthesis (Eurofins Genomics) and cloned into the HindIII/XhoI sites of pcDNA3.1. The expression vector encoding eIF4G1 with deletion of the HEAT3 domain (4G1-ΔHEAT3 [aa 192-1437]) was generated from the long HA-eIF4G1 vector by inserting a stop codon (nt 4312-4313, GC to TG) by site-directed mutagenesis (Q5 kit, New England Biolabs). All plasmids were amplified into *E. coli* DH5α, and insert sequences were validated by sequencing.

### Expression of human recombinant FKBP7

For recombinant protein production, purified pET28b(+) plasmids encoding the FKBP7 PPIase domain (aa 31-145) were transformed by heat shock into *E. coli* BL21* (DE3) (ThermoFisher). Cells were grown in M9 medium supplemented with $^{15}$N-ammonium chloride at 37°C until the optical density at 600 nm reached 0.6-0.8, then induced with 0.5 mM isopropyl β-D-1-thiogalactopyranoside (IPTG) and incubated overnight at 20°C. Cells were harvested by centrifugation at 6000 rpm for 30 min at 4 °C, and the pellet was frozen at -80°C until purification.

**Purification of human recombinant FKBP7**

Cell pellets were resuspended in Buffer A (50 mM Tris pH 8, 200 mM NaCl, 2 mM DTT) supplemented with EDTA-free protease inhibitor cocktail (Roche) and lysozyme (10mg per pellet), then passed three times through a French press at 1.5 kbar to lyse the cells. To isolate the soluble fraction from cellular debris, the lysate was centrifuged at 100,000xg for 1 h at 4°C. The protein solution was filtered through a 0.45 µm membrane and loaded onto a $Ni^{2+}$-chelating column (GE Healthcare, HisTrap FF crude). After washing with buffer A and removing weakly bound proteins with buffer A containing 10 mM and 60 mM imidazole, FKBP7 was eluted with elution buffer (20 mM Tris pH 8, 200 mM NaCl, 2 mM DTT, 300 mM imidazole). Fractions containing the protein were pooled and concentrated with a 10 kDa centrifugal filter unit, then subjected to gel filtration. The protein solution was diluted in 10 ml of 50 mM HEPES pH 7, 150 mM NaCl, 2 mM TCEP, concentrated to a final volume of 500 µl, and injected onto Superdex 75 10/300 GL Column (Sigma-Aldrich). His6-tag was removed by overnight incubation with homemade TEV protease at 25°C (1:40 TEV:protein mass ratio). The protein solution was then diluted 2.5-fold with 50 mM HEPES pH 9, 150 mM NaCl, 2 mM TCEP to raise the pH to 8 and increase His6-tag binding to a Ni-NTA column. The resulting solution was incubated with HisPur Ni-NTA resin (ThermoFisher) for 30 min at 4°C to remove the cleaved-tag, uncleaved protein, and the TEV protease, and the soluble fraction was collected by gravity-flow chromatography (Bio-Rad) and purified with a HiPrep 26/10 desalting column (Sigma-Aldrich) into final buffer (50 mM HEPES pH 7, 150 mM NaCl, 2 mM TCEP, 2 mM EDTA). Protein fractions were pooled and frozen with liquid nitrogen. The quality of recombinant FKBP7 was assessed by SDS-PAGE and Coomassie blue staining.

**Bioinformatic analysis**

Fastq files were trimmed and quality-checked with Fastp *(69)*, trimming both ends with a sliding window of size 6 and a minimum quality of 10. Reads shorter than 15 bases were discarded, reads with more than 50 % unqualified bases were removed, and over representation analysis was enabled. Transcript abundance was estimated using GRCh38.99 Ensembl genome with Salmon *(70)*. A decoy-aware gentrome index was built while keeping transcripts with identical sequences and different genomic loci. Pseudo-mapping was performed with correction for GC content, sequence bias awareness and positional bias consideration, since the prior trimming step was expected RNA sequences bias from sequencing. One hundred bootstraps were performed to assess sequencing errors. Transcript counts were aggregated to gene counts with tximport *(71)*, and only genes with an expression level of at least 50 in every sample were retained for downstream analysis. Anota2seq *(41)* was used with default parameters to identify statistically significant changes in translation, and batch effect between replicates was corrected with batchVec parameter of the anota2seqDataSetFromMatrix function. Analysis quality was assessed with Samtools *(72)* and RSeQC *(73)*, and FastQ Screen *(74)* was used to test for sample contamination, returning a clean profile. Quality reports were aggregated with MultiQC *(75)*. The counts-table generation pipeline was built with Snakemake *(76)*. Translation-change analysis was performed in R 4.3.1 within a Singularity container. Scripts and pipeline are available on Zenodo. Gene set enrichment analysis (GSEA) was performed using the 2,085 unique buffering gene identified by anota2seq. Genes were ranked according to the log2 fold change and analyzed using the GSEA Preranked tool (v4.3.3, Broad Institute). Enrichment was assessed against the Reactome and WikiPathways gene set collections. Gene sets with a false discovery rate (FDR) q-value < 0.05 were considered significantly enriched, then ranked by normalized enrichment score (NES).
