## Supplemental figures and legends for "Cytoplasmic FKBP7 reflux sustains NFE2L1 levels in an adaptive response to chemotherapy in prostate cancer cells"

**Supplemental Figure 1**

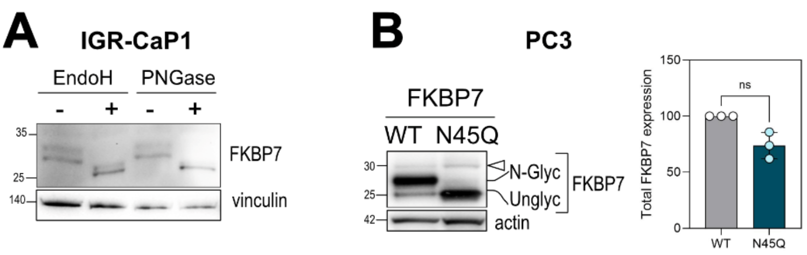


**Figure S1: FKBP7 is N-glycosylated.**

**A.** Immunoblot showing FKBP7 N-glycosylation status in lysates from IGR-CaP1 cells left untreated or treated with Endoglycosidase H or PNGase F. Vinculin is the loading control. **B.** Immunoblot showing the identification of the FKBP7 N-glycosylation site after transfection of an expression vector encoding wild-type FKBP7 (WT) or the N45Q point mutant in PC3 cells. Bands are labeled for the N-glycosylated (N-Glyc) and non-glycosylated (Unglyc) forms. The white triangle indicates the additional FKBP7 band at 30 kDa, visible for both WT and the N45Q mutant. Actin is the loading control. Quantification of WT FKBP7 and the N45Q mutant form in total extracts is shown on the right (from three independent experiments, ratio paired t test).

**Supplemental Figure 2**

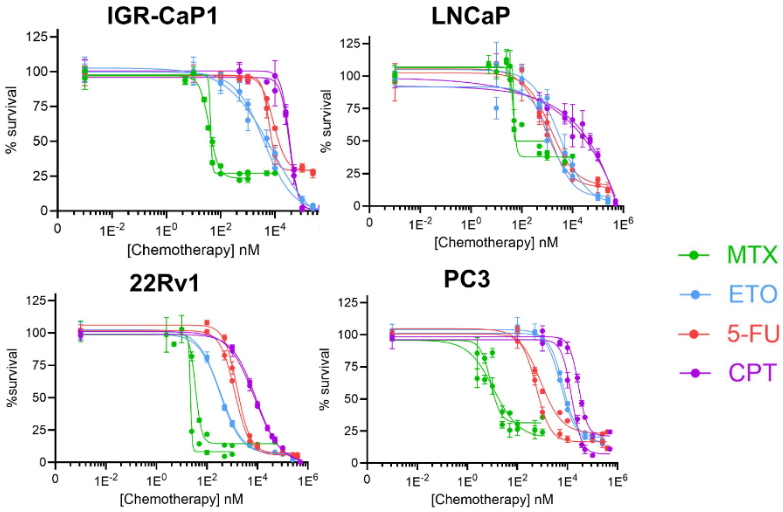


| | IGR-CaP1 | LNCaP | 22Rv1 | PC3 |
|---|---|---|---|---|
| Chemotherapy | | Mean IC50 | | |
| MTX | 41 nM | 47 nM | 29 nM | 10 nM |
| ETO | 6 µM | 2.4 µM | 338 nM | 6 µM |
| 5-FU | 8 µM | 1.1 µM | 1.5 µM | 709 nM |
| CPT | 35 µM | ~50 µM | 8.3 µM | 21 µM |

**Figure S2: Chemotherapy dose–response and $IC_{50}$ in prostate cancer cells.**

Viability was quantified in four prostate cancer cell lines after 72h incubation with methotrexate (MTX), etoposide (ETO), 5-fluorouracil (5-FU) or carboplatin (CPT). Data points are mean ± SD of two independent experiments. $IC_{50}$ values were calculated by GraphPad Prism software and are expressed as means obtained from two independent determinations.

**Supplemental Figure 3**

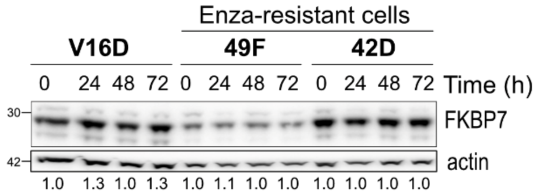


**Figure S3: AR-targeted therapy does not increase FKBP7 levels.**

Immunoblots showing total FKBP7 levels in response to treatment with 10µM enzalutamide in the LNCaP-derived V16D (parental) or enzalutamide-resistant (49F and 42D) cells *(61)*. Actin is the loading control. Protein levels were quantified using Image Lab software. FKBP7 is expressed relative to the corresponding loading control and quantification relative to untreated V16D cells is shown below the image.

**Supplemental Figure 4**

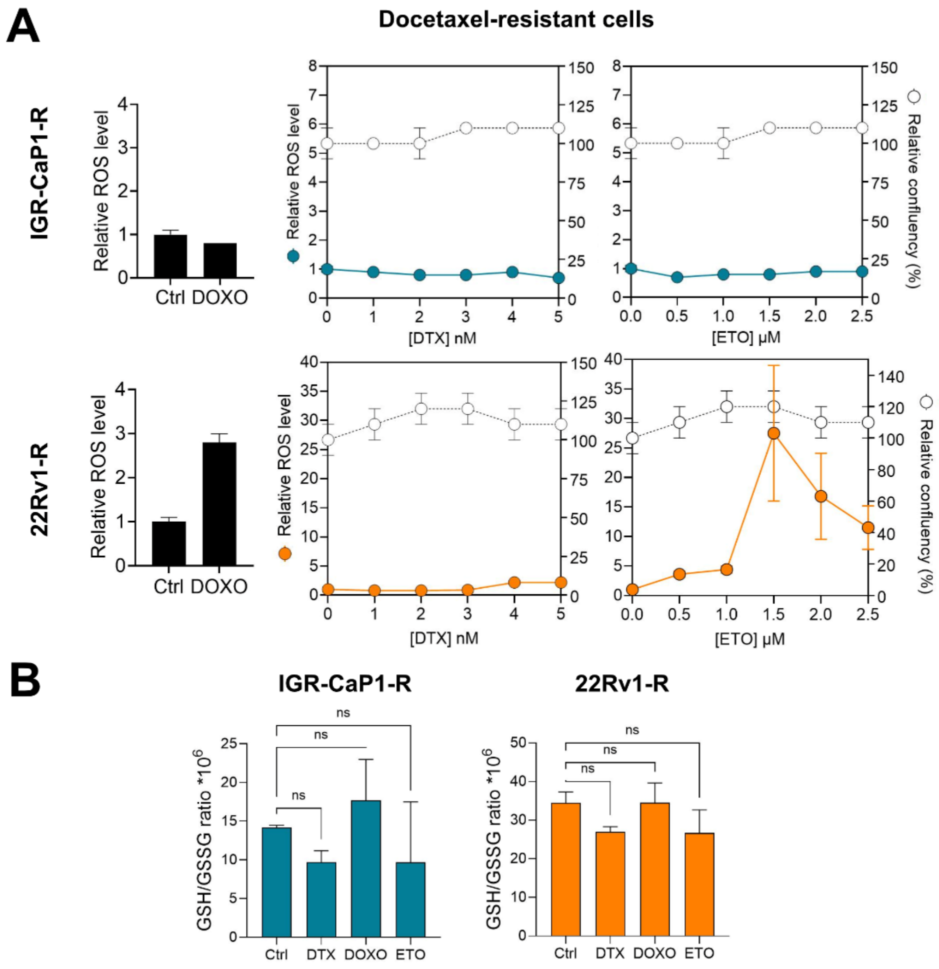


**Figure S4: Effect of oxidative stressors on ROS content in parental and resistant cells.**

**A.** ROS levels, quantified using $H_2DCFDA$, and cell confluence were simultaneously determined after IGR-CaP1 and 22Rv1 docetaxel-resistant cells were treated for 48 h with 100 nM Doxorubicin (DOXO), or with increasing doses of docetaxel (DTX) and etoposide (ETO). ROS levels were expressed as Integrated Intensity Average Object/Area confluence. The curves report both relative ROS levels (color dots) and cell confluence (empty dots). The DOXO condition was represented on a histogram as control. Data normalized to control Mock condition are mean ± SEM. **B.** Determination of GSH/GSSG ratio after 48 h of treatment of the resistant cells with 125 nM DTX, DOXO (100 nM in IGR-CaP1-R and 10 nM in 22Rv1-R cells) or ETO (5 µM in resistant IGR-CaP1-R and 1 µM in resistant 22Rv1-R cells). Data representative of two independent experiments, are mean ± SD. ns, non-significant as determined by one-way ANOVA with Dunnett's post-hoc test.

**Supplemental Figure 5**

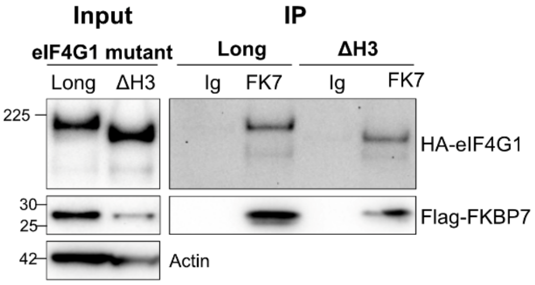


**Figure S5: FKBP7 binds eIF4G1 ΔH3 mutant deleted of the C-terminal HEAT3 domain.**
FKBP7 was immunoprecipitated with the anti-FKBP7 antibody (or IgG as control) in PC3 cells co-transfected with Flag-FKBP7-WT and HA-eIF4G1 (4G1-long or ΔHEAT3) mutant encoding vectors. Immunoblot showing FKBP7 and the co-immunoprecipitated HA-eIF4G1 mutants (Long and ΔH3), revealed with the anti-HA antibody. FKBP7 was revealed with anti-Flag antibody. Input controls (10%) are shown.

**Supplemental Figure 6**

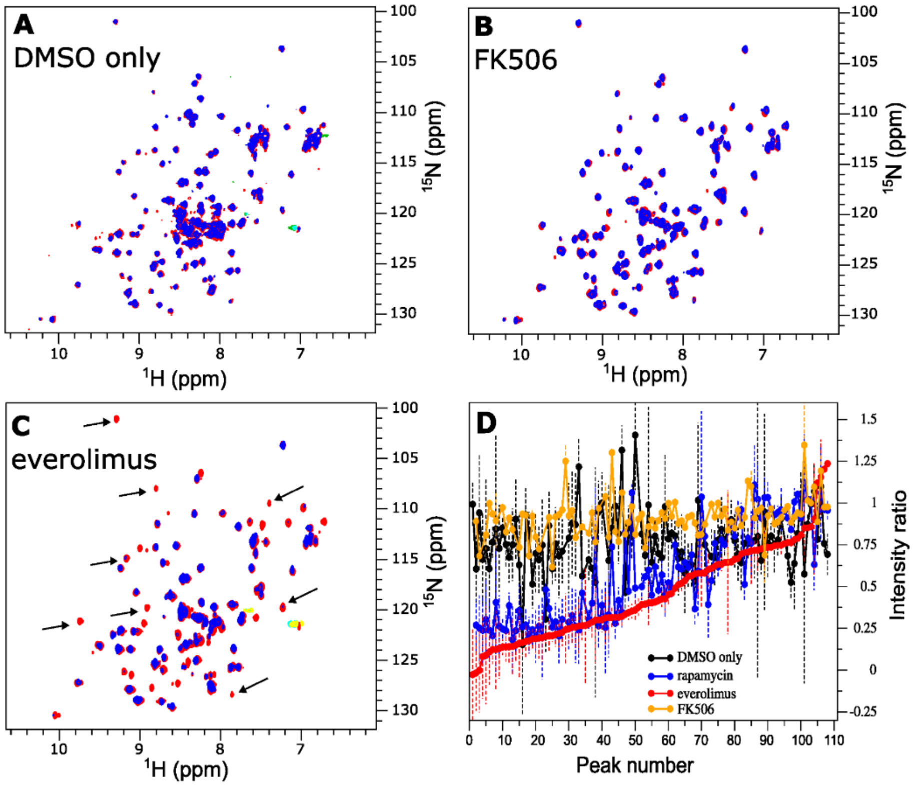


**Figure S6: FKBP7 can bind everolimus but not FK506.**

$^{15}$N SOFAST-HMQC NMR spectra of $^{15}$N-labeled FKBP7 PPI domain without (blue) and with (red) DMSO (**A**), 1:2 molar ratio FK506 (**B**), or 1:1 molar ratio everolimus (**C**). All ligand stocks were prepared in 100 % DMSO. Each crosspeak corresponds to one H-N chemical bond, corresponding to approximately one amino acid residue. Most, but not all, crosspeaks had reduced intensity upon addition of everolimus, and some (labeled by an arrow) completely disappeared. **D.** Intensity loss upon DMSO or ligand/DMSO addition calculated from peak intensity ratios extracted from the NMR spectra. In the absence of resonance assignments, the peaks were ordered according to the increasing intensity ratio values obtained for everolimus (red curve). Only minor, non-specific intensity loss was observed upon addition of DMSO only or with FK506/DMSO while a subset of peaks showed stronger intensity reduction upon addition of rapamycin and everolimus. The plot indicates that the same amino-acid residues underwent strong signal reduction upon addition of everolimus or rapamycin, suggesting that the two ligands occupy the same binding site in FKBP7.

**Supplemental Figure 7**

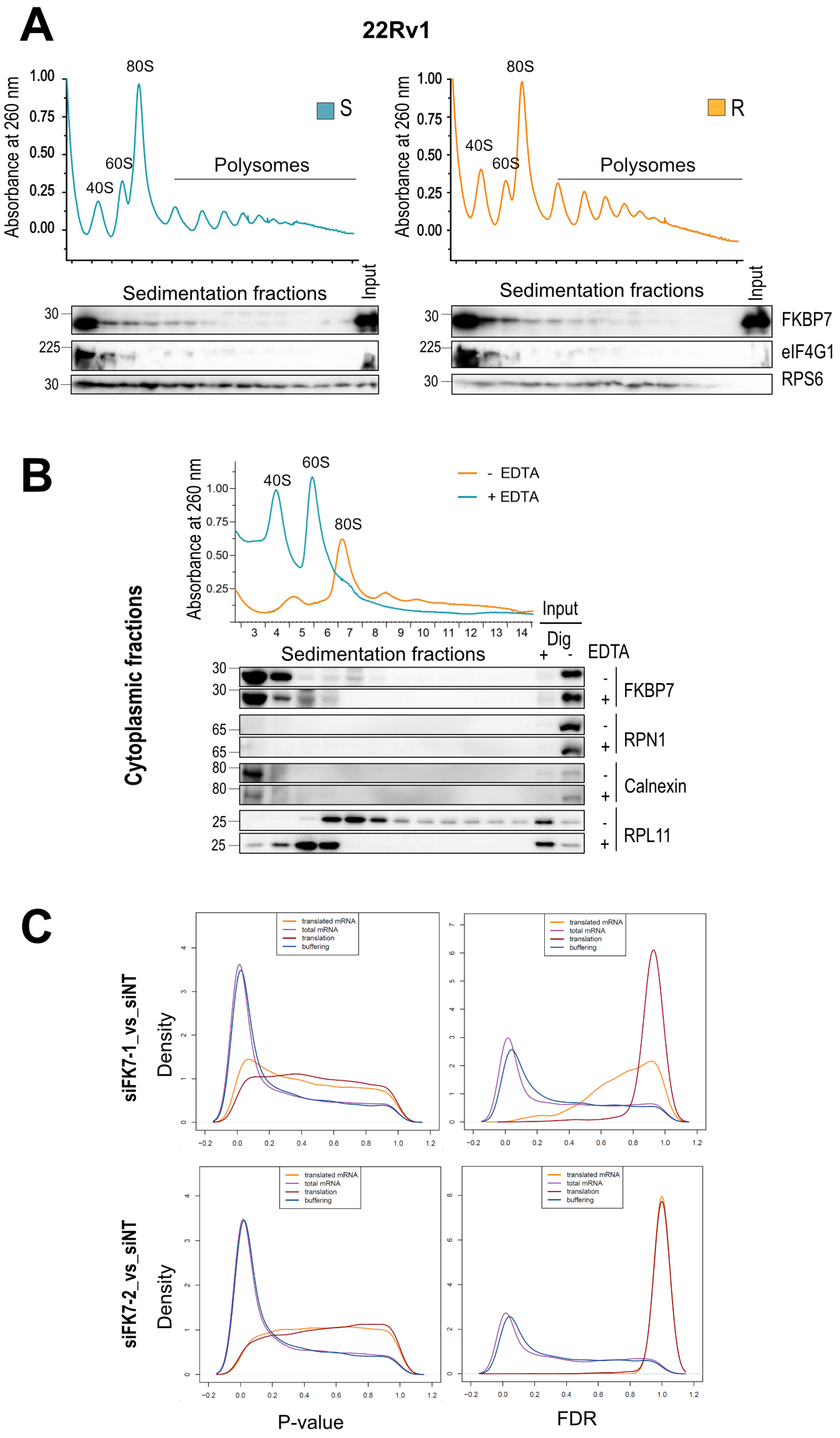

**Figure S7: FKBP7 interacts with eIF4G1 and the 40S ribosomal subunit and contributes to translation. A.** Polysome profiles of parental (blue) and docetaxel-resistant (orange) 22Rv1 cell lines under basal conditions (representative of three independent experiments). Immunoblot showing FKBP7, eIF4G1, and RPS6 (marker for the 40S ribosomal subunit) in each fraction. Input corresponds to the cellular extract before loading onto sucrose gradient. **B.** Cytosolic ribosomal complexes were isolated from docetaxel-resistant 22Rv1 cells using a digitonin-based permeabilization protocol to exclude ER-derived material. Input lysates were kept before (dig-) and after digitonin permeabilization (dig+, 1/100 of the total extracts). The resulting cytosolic ribosomal fraction was left untreated (-EDTA, orange) or treated with EDTA (+EDTA, blue), and both samples were resolved on 5-50% sucrose gradients monitored by absorbance at 260 nm (top). Input and gradient fractions were analyzed by western blot for FKBP7, RPN1 and calnexin as ER markers, and RPL11 for 60S ribosomal subunit. **C.** Distributions (quantified by kernel density estimation) of P-values and FDR-adjusted P-values for differential expression siFK1_vs_siNT and siFK7-2_vs_siNT, in resistant IGR-CaP1 cells using data from polysome-associated mRNA (orange) or cytosolic mRNA (purple), for analysis of differences in translational efficiencies leading to altered protein levels (red), and analysis of translational buffering (blue).

### Supplemental Figure 8

**A**

**WIKI canonical pathways (UP)**

| NAME | SIZE | ES | NES | NOM p-val | FDR q-val |
|---|---|---|---|---|---|
| WP_CHOLESTEROL_BIOSYNTHESIS_PATHWAY_IN_HEPATOCYTES | 15 | 0,7032 | 2,2465322 | 0,0000000 | 0,0000000 |
| WP_FATTY_ACID_AND_LIPOPROTEIN_TRANSPORT_IN_HEPATOCYTES | 29 | 0,4655 | 1,7859863 | 0,0072464 | 0,0440700 |
| WP_NRF2_PATHWAY | 18 | 0,5328 | 1,7468591 | 0,0098684 | 0,0385149 |
| WP_NUCLEAR_RECEPTORS_METAPATHWAY | 33 | 0,4086 | 1,5830986 | 0,0248963 | 0,0919339 |
| WP_EBOLA_VIRUS_INFECTION_IN_HOST | 15 | 0,3933 | 1,2709311 | 0,1843750 | 0,3810906 |
| WP_CYTOPLASMIC_RIBOSOMAL_PROTEINS | 36 | 0,3162 | 1,2616987 | 0,1275720 | 0,3313336 |

**WIKI canonical pathways (DOWN)**

| NAME | SIZE | ES | NES | NOM p-val | FDR q-val |
|---|---|---|---|---|---|
| WP_RETINOBLASTOMA_GENE_IN_CANCER | 29 | -0,7358 | -2,3782716 | 0 | 0 |
| WP_DNA_REPAIR_PATHWAYS_FULL_NETWORK | 32 | -0,6824 | -2,1975887 | 0 | 0 |
| WP_DNA_REPLICATION | 15 | -0,8074 | -2,1924057 | 0 | 0 |
| WP_PRIMARY_OVARIAN_INSUFFICIENCY | 17 | -0,7756 | -2,1317410 | 0 | 0 |
| WP_DNA_IRDAMAGE_AND_CELLULAR_RESPONSE_VIA_ATR | 18 | -0,7349 | -2,0807247 | 0 | 2,E-04 |
| WP_CELL_CYCLE | 24 | -0,6002 | -1,8528149 | 0,0056180 | 0,0120423 |

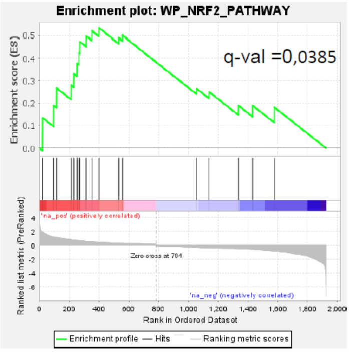

B

Up-regulated Pathways

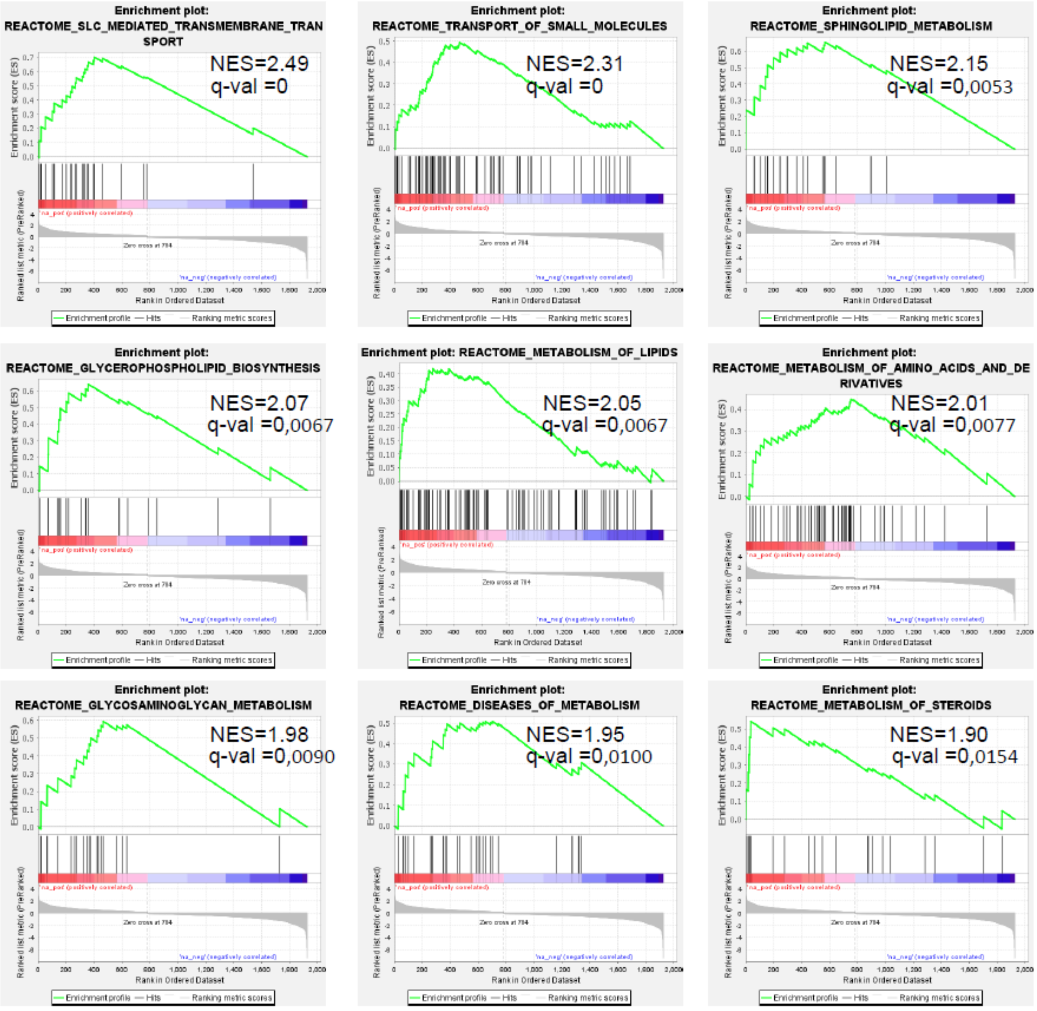

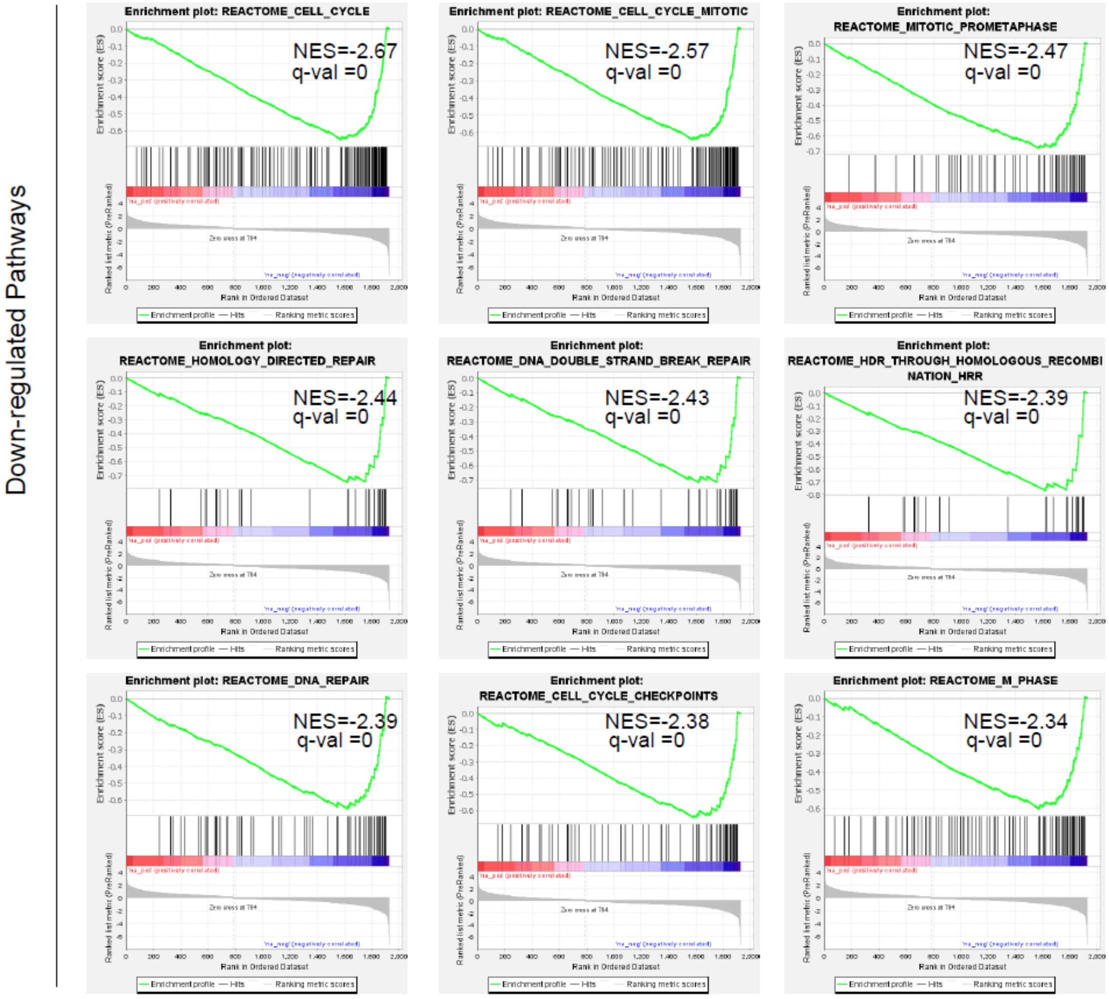


**Figure S8: FKBP7 silencing-mediated translational buffering modulates the expression of target genes of NFE2L1 and NFE2L2 transcription factors.**

**A.** Gene Set Enrichment Analysis (GSEA) on the buffering gene set (2,085 genes) derived from anota2seq analysis. Left, panel display top six normalized enrichment scores (NES) for the up-regulated and down-regulated WIKI canonical pathways, respectively. Right, enrichment diagram of the NFE2L2/NRF2 WIKI pathway. **B.** The top nine ranking diagrams of the REACTOME pathways from the GSEA analysis, with NES and FDR q-values were selected from among the 13 significantly enriched up-regulated and 56 down-regulated pathways (FDR < 0.05).

### Supplemental Figure 9

**A**

| mRNA Regulation mode | Total mRNA | NFE2L1 targets | NFE2L2 targets | NFE2L3 targets | All NRF targets |
|---|---|---|---|---|---|
| Buffering | 1391 | 503 | 213 | 57 | 56% |
| Abundance | 115 | 39 | 10 | 2 | 44% |

**B**

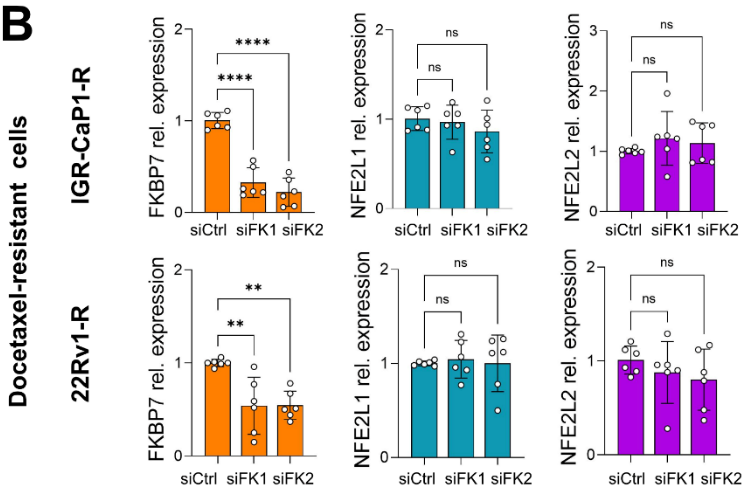


**Figure S9: FKBP7 silencing does not decrease NFE2L1 or NFE2L2 transcriptional expression in resistant cells.**
**A.** Number of genes corresponding to NFE2L1, NFE2L2, and NFE2L3 targets associated with the Liu et al. 2019 signature *(44)*, found in the translational buffering gene and abundance gene sets, respectively. **B.** Quantitative RT-PCR of FKBP7, NFE2L1 and NFE2L2 mRNA in docetaxel-resistant IGR-CaP1 and 22Rv1 cells transfected with siNT or two siRNA targeting FKBP7. The gene levels were normalized to siNT condition. Data (from three independent experiments) represent mean ± SD. *P values* were obtained using one-way ANOVA with Dunnett's posttest (**p <0.01; ***p <0.001; ns : non-significant).

### Supplemental Figure 10

```
FKBP12 Numbering        2      10          20        30       36       40         50          60        70        80         90       99     107
P62942|FKB1A      3   --VQVETISPGD-GRTFP-KRGQTCVVHYTGMLED---GKKFD-----SSRDRN--KPFKFMLGKQEVIR---GWEEGVAQMSVGQRAKLTISPDYAYGATG-HPGIIPPHATLVFDVELLKLE----
P68106|FKB1B      3   --VEIETISPGD-GRTFP-KKGQTCVVHYTGMLQN---GKKFD-----SSRDRN--KPFKFRIGKQEVIK---GFEEGAAQMSLGQRAKLTCTPDVAYGATG-HPGVIPPNATLIFDVELLNLE----
O75344|FKBP6      38  --VLKDVIREGA-GDLVA--PDASVLVKYSGYLEHM--DRPFD-----SNYFRK--TPRLMKLGEDITLW---GMELGLLSMRRGELARFLFKPNYAYGTLG-CPPLIPPNTTVLFEIELLDFLDC--
Q14318|FKBP8      102 --LRKKTLVPGPPGSSRP-VKGQVVTVHLQTSLEN---GTRVQ-----EE------PELVFTLGDCDVIQ---ALDLSVPLMDVGETAMVTADSKYCYGPQG-RSPYIPPHAALCLEVTLKTAVDG--
Q02790|FKBP4_FK1  33  --VLKVIKREGT-GTEMP-MIGDRVFVHYTGWLLD---GTKFD-----SSLDRK--DKFSFDLGKGEVIK---AWDIAIATMKVGEVCHITCKPEYAYGSAG-SPPKIPPNATLVFEVELFEFKGE--
Q02790|FKBP4_FK2  150 --IIRRIQTRGE-GYAKP-NEGAIVEVALEGYYK----DKLFD-----Q-------RELRFEIGEGENLDLPYGLERAIQRMEKGEHSIVYLKPSYAFGSVGKEKFQIPPNAELKYELHLKSFEKA--
Q13451|FKBP5_FK1  33  --VLKIVKRVGN-GEETP-MIGDKVYVHYKGKLSN---GKKFD-----SSHDRN--EPFVFSLGKGQVIK---AWDIGVATMKKGEICHLLCKPEYAYGSAG-SLPKIPSNATLFFEIELLDFKGE--
Q13451|FKBP5_FK2  148 --IIRRTKRKGE-GYSNP-NEGATVEIHLEGRCG----GRMFD-----C-------RDVAFTVGEGEDHDIPIGIDKALEKMQREEQCILYLGPRYGFGEAG-KPKFGIEPNAELIYEVTLKSFEKA--
Q00688|FKBP3      111 --YTKSVLKKGD-KTNFP-KKGDVVHCWYTGTLQD---GTVFDTNIQTSAKKKKNAKPLSFKVGVGKVIR---GWDEALLTMSKGEKARLEIEPEWAYGKKGQPDAKIPPNAKLTFEVELVDID----
Q5T1M5|FKB15      181 --VLSQDLIVAD-GPAVE-VGDSLEVAYTGWLFQNHVLGQVFD-----STANKD--KLLRLKLGSGKVIK---GWEDGMLGMKKGGKRLLIVPPACAVGSEGVIGWTQATDSILVFEVEVRRVKFA--
P26885|FKBP2      30  KLQIGVKKRVDH-CPIKS-RKGDVLHMHYTGKLE-D---GTEFD-----SSLPQN--QPFVFSLGTGQVIK---GWDQGLLGMCEGEKRKLVIPSELGYGERG-APPKIPGGATLVFEVELLKIER---
Q9NYL4|FKB11      39  --QVETLVEPPEPCAEPA-AFGDTLHIHYTGSLV-D---GRIID-----TSLTR---DPLVIELGQKQVIP---GLEQSLLDMCVGEKRRAIIPSHLAYGKRG-FPPSVPADAVVQYDVELIALIRA--
Q9NWM8|FKB14      28  --KIEVLQKPFI-CHRKT-KGGDLMLVHYEGYLEKD---GSLFH-----STHKHNNGQPIWFTLGILEALK---GWDQGLKGMCVGEKRKLIIPPALGYGKEG-KGK-IPPESTLIFNIDLLEIRNG--
Q9Y680|FKBP7      36  --KIEVLHRPEN-CSKTS-KKGDLLNAHYDGYLAKD---GSKFY-----CSRTQNEGHPKWFVLGVGQVIK---GLDIAMTDMCPGEKRKVVIPPSFAYGKEGYAEGKIPPDATLIFEIELYAVTKG--
O95302|FKBP9_FK1  43  --QIERRFVPDE-CPRTV-RSGDFVRYHYVGTFP-D---GQKFD-----SSYDRD--STFNVFVGKGQLIT---GMDQALVGMCVNERRFVKIPPKLAYGNEG-VSGVIPPNSVLHFDVLLMDIWN---
O95302|FKBP9_FK2  149 --QIHTYFKPPS-CPRTI-QVSDFVRYHYNGTFL-D---GTLFD-----SSHNRM--KTYDTYVGIGWLIP---GMDKGLLGMCVGEKRIITIPPFLAYGEDG-DGKDIPGQASLVFDVALLDLH----
O95302|FKBP9_FK3  262 ---IENKVVPEN-CERIS-QSGDFLRYHYNGTLL-D---GTLFD-----SSYSRN--RTFDTYIGQGYVIP---GMDEGLLGVCIGEKRRIVVPPHLGYGEEG-RGN-IPGSAVLVFDIHVIDFHN---
O95302|FKBP9_FK4  372 --SITSHYKPPD-CSVLS-KKGDYLKYHYNASLL-D---GTLLD-----STWNLG--KTYNIVLGSGQVVL---GMDMGLREMCVGEKRTVIIPPHLGYGEAG-VDGEVPGSAVLVFDIELLELVAG--
Q96AY3|FKB10_FK1  44  -VVIERYHIPRA-CPREV-QMGDFVRYHYNGTFE-D---GKKFD-----SSYDRN--TLVAIVVGVGRLIT---GMDRGLMGMCVNERRRLIVPPHLGYGSIG-LAGLIPPDATLYFDVVLLDVWNK--
Q96AY3|FKB10_FK2  163 --QVSTLLRPPH-CPRMV-QDGDFVRYHYNGTLL-D---GTSFD-----TSYSKG--GTYDTYVGSGWLIK---GMDQGLLGMCPGERRKIIIPPFLAYGEKG-YGTVIPPQASLVFHVLLIDVHN---
Q96AY3|FKB10_FK3  269 --QLETLELPPG-CVRRA-GAGDFMRYHYNGSLM-D---GTLFD-----SSYSRN--HTYNTYIGQGYIIP---GMDQGLQGACMGERRRITIPPHLAYGENG-TGDKIPGSAVLIFNVHVIDFHN---
Q96AY3|FKB10_FK4  383 ----RTLSRPSETCNETT-KLGDFVRYHYNCSLL-D---GTQLF-----TSHDYG--APQEATLGANKVIE---GLDTGLQGMCVGERRQLIVPPHLAHGESG-ARG-VPGSAVLLFEVELVSRED---
FKBP7 numbering                                          71                                                                           137
```

**Figure S10: Conservation of Phe amino acids in the PPIase domain of FKBP7 among FKBPs.**
Alignment of primary sequences of the FKBP PPIase domains showing high conservation of residues Phe36 and Phe99 (FKBP12 numbering), which correspond to Phe71 and Phe137 in FKBP7 and are critical for isomerase activity. Red, FKBP12 numbering; blue, FKBP7 numbering.
