## Supplemental Table 1 for "Cytoplasmic FKBP7 reflux sustains NFE2L1 levels in an adaptive response to chemotherapy in prostate cancer cells"

**Supplementary Table 1**

| REAGENT or RESOURCE | SOURCE | IDENTIFIER |
|---|---|---|
| **Antibodies** | | |
| Rabbit polyclonal anti-FKBP7 | Protein Atlas | Cat#HPA008707; RRID:AB_1848624 |
| Normal rabbit anti-IgG | Cell Signaling Technology | Cat# 2729 RRID:AB_1031062 |
| Rabbit polyclonal anti-GFP | AbCam | Cat# Ab290 RRID:AB_303395 |
| Rabbit monoclonal anti-eIF4G1 (D6A6) | Cell Signaling Technology | Cat# 8701 RRID:AB_11178378 |
| Rabbit monoclonal anti-HA-tag (C29F4) | Cell Signaling Technology | Cat# 3724 RRID:AB_1549585 |
| Rabbit monoclonal anti-IRE1α (14C10) | Cell Signaling Technology | Cat# 3294 RRID:AB_823545 |
| Rabbit monoclonal anti-PERK (C33E10) | Cell Signaling Technology | Cat# 3192 RRID:AB_2095847 |
| Rabbit monoclonal anti-p21 Waf1/Cip1 (12D1) | Cell Signaling Technology | Cat# 2947 RRID:AB_823586 |
| Rabbit monoclonal anti-vinculin (E1E9V) | Cell Signaling Technology | Cat# 13901 RRID:AB_2728768 |
| Rabbit polyclonal anti-NFE2L1 | Proteintech | Cat# 1702-1-AP RRID:AB_2878342 |
| Rabbit polyclonal anti-RPL11 | Proteintech | Cat# 16277-1-AP RRID:AB_2181292 |
| Rabbit polyclonal anti-RPN1 | Proteintech | Cat# 12894-1-AP RRID:AB_2170474 |
| Mouse monoclonal anti-HA-tag (6E2) HRP-conjugate | Cell Signaling Technology | Cat# 2999 RRID:AB_1264166 |
| Mouse monoclonal anti-S6 ribosomal protein (RPS6) (54D2) | Cell Signaling Technology | Cat# 2317 RRID:AB_2238583 |
| Mouse monoclonal anti-Flag-tag (2B3C4) | Proteintech | Cat# 66008-3-Ig RRID:AB_2749837 |
| Mouse monoclonal anti-ATF6 (3B7E4) | Proteintech | Cat# 66563-1-Ig RRID:AB_2881924 |
| Mouse monoclonal anti-NFE2L2 (A-10) | Santa-Cruz Biotechnology | Cat# sc-365949 RRID:AB_10917561 |
| Mouse monoclonal anti-HSC70 (B-6) | Santa-Cruz Biotechnology | Cat# sc-7298 RRID:AB_627761 |
| Mouse monoclonal anti-calnexin (E-10) | Santa-Cruz Biotechnology | Cat# sc-46669 RRID:AB_626784 |

| | | |
|---|---|---|
| Mouse monoclonal anti- β-actin (AC-15) | Sigma-Aldrich | Cat# A5441 RRID:AB_476744 |
| Mouse monoclonal anti-α-tubulin (DM1A) | Sigma-Aldrich | Cat# T9026 RRID:AB_477593 |
| **siRNA Oligonucleotides** | | |
| Stealth-siRNA targeting sequence: FKBP7 #1: GGCCUAGACAUUGCUAUGACAGAUA | Thermo Fisher Scientific | Cat# 10620318 (HSS122495) |
| Stealth-siRNA targeting sequence: FKBP7 #2: CCUCUACUUGCAAAGGGAAUUUGAA | Thermo Fisher Scientific | Cat# 10620318 (HSS182176) |
| Silencer select siRNA targeting sequence: FKBP7 #1: GGCUCGAAAUUCUACUGCAtt | Thermo Fisher Scientific | Cat# 4390824 (s28491) |
| Silencer select siRNA targeting sequence: FKBP7 #2: GACAUUGCUAUGACAGAUAtt | Thermo Fisher Scientific | Cat# 4390824 (s28492 |
| Silencer select siRNA targeting sequence: NFE2L1#1: GCUGCGAGAUGAGAACGGAtt | Thermo Fisher Scientific | Cat# 4392420 (s9488) |
| Silencer select siRNA targeting sequence: NFE2L1#2: GGGUGGACGUGGAUACUUAtt | Thermo Fisher Scientific | Cat# 4392420 (s9490) |
| Silencer select siRNA targeting sequence: NFE2L1#3: GGCGUGAGGUUUUUGACUAtt | Thermo Fisher Scientific | Cat# 4392420 (s9489) |
| Stealth RNAi™ siRNA Negative Control | Thermo Fisher Scientific | Cat# 12935300 |
| Silencer™ Select Negative Control | Thermo Fisher Scientific | Cat# 4390846 |
| **PCR primers** | | |
| PCR primer: FW-FKBP7: 5’-GCATCGTCCAGAAAACTGCT-3’ | Eurofins Genomics | |
| PCR primer: FW-NFE2L1: 5’-AGTGGAGACTTAACCAAAGAGGAC-3’ | Eurofins Genomics | |
| PCR primer: FW-NFE2L2: 5’-TCCAGTCAGAAACCAGTGGAT-3’ | Eurofins Genomics | |
| PCR primer: FW-TBP: 5’-CCCATGACTCCCATGACC-3’ | Eurofins Genomics | |

Antibodies, siRNA and PCR primers used in this study
